# Strong GAL4 expression compromises *Drosophila* fat body function

**DOI:** 10.1101/2025.07.21.665961

**Authors:** Scott A. Keith, Ananda A. Kalukin, Dana S. Vargas Solivan, Melanie R. Smee, Brian P. Lazzaro

**Author notes:** Corresponding author: Scott A. Keith.

## Abstract

The ability to direct tissue-specific overexpression of transgenic proteins in genetically tractable organisms like *Drosophila melanogaster* has facilitated innumerable biological discoveries. However, transgenic proteins can themselves impact cellular and physiological processes in ways that are often ignored or poorly defined. Here we discovered that the *yolk-GAL4* transgene, which directs strong expression of the yeast GAL4 transcription factor in the *Drosophila* fat body, induces significant physiological defects in adult female flies. We found that *yolk-GAL4* disrupts adipose tissue integrity and reduces fat body lipid stores, egg production, and resistance to systemic bacterial infections. Knocking down *GAL4* expression in *yolk-GAL4* heterozygotes using RNAi fully suppressed each of these defects, thus confirming that the GAL4 transgene product induces these phenotypes. Comparing a panel of additional fat body driver lines, we found that *GAL4* expression levels directly correlate infection susceptibility, but not with fat levels or egg production. To determine whether other transgenic proteins can impair fat body function, we constructed new fly lines in which the *yolk* enhancer directs expression of either cytoplasmic or nuclear-localized mCherry, or an alternative transactivator, LexA. We found that only nuclear-localized mCherry and LexA increased infection susceptibility similarly to GAL4, suggesting that intranuclear transgenic proteins in general can curtail the fat body’s induced immune response in a manner highly sensitive to transgene expression strength. Additionally, these new lines can be valuable tools for future studies. More broadly, our findings highlight the potential for transgenes to substantially impact organismal biology and emphasize the importance of rigorously characterizing genetic tools to optimally leverage model systems like *Drosophila*.

## INTRODUCTION

Cells and tissues dynamically modulate transcription, translation, metabolism, secretion, and other functions to sustain the health and fitness of multicellular organisms. Internal and external factors that disrupt the tightly coordinated balance among these processes, or that overburden any particular process, can induce cellular strain leading to tissue– and organism-level dysfunction. In some cases, the introduction of transgenes utilized to study gene expression and protein function can disrupt tissue homeostasis in genetically amenable model organisms (Liu et al. 1999; Huang et al. 2000; Detrait et al. 2002; Kramer and Staveley 2003; Krestel et al. 2004; Rezával et al. 2007; Liu and Lehmann 2008; Kintaka et al. 2016; Kintaka et al. 2020; Namba and Moriya 2024; Verma et al. 2024). These effects can complicate molecular investigations because high-level expression of the transgene is often desired to maximize the strength of experimental manipulations. However, overly strong expression and consequent high levels of the transgenic protein product may induce cellular strain that inadvertently affects the traits being studied, potentially confounding observed phenotypes and their biological significance.

The advent of the *GAL4-UAS* system marked a watershed advance for genetic experimentation in *Drosophila melanogaster* (hereafter referred to as “*Drosophila*”). In this bipartite expression system, *Drosophila* promoter or enhancer sequences direct expression of the *Saccharomyces cerevisiae* GAL4 transcription factor, either ubiquitously or in specific tissues and/or developmental stages. Flies expressing GAL4 under the desired regulation are then crossed to flies carrying a transgene downstream of the yeast *Upstream Activation Sequence* (*UAS*), which is a transcriptional promoter recognized by GAL4. Resultant F1 progeny bearing both genetic elements will thus express the *UAS*-linked transgene only in the cells or tissues producing GAL4 (Brand and Perrimon 1993). This system thereby enables facile, precise, and sophisticated genetic manipulations via simple experimental mating schemes. The *GAL4-UAS* expression system is extremely widely used, including for spatially-localized induction of RNA hairpin constructs that reduce expression levels of individual genes by RNA interference (RNAi), overproduction or ectopic expression of either native or heterologous proteins, and for visually labeling cells and subcellular structures with fluorescent or epitope-recognizable molecules (reviewed in Caygill and Brand 2016). The versatility of this system and the vast collection of extant driver and responder lines have facilitated innumerable biological discoveries in *Drosophila*. However, the potential effects of GAL4 itself on fly cell biology and physiology are rarely considered and often remain poorly defined (but see Kramer et al. 2003; Rezával et al. 2007; Liu and Lehmann 2008; Zappia et al. 2024).

The fat body is a tissue that governs whole-organism physiology in *Drosophila* and other insects. Composed of segmented, planar sheets of adipocytes, the fat body is distributed throughout the body segments, lining the cuticle and surrounding the visceral organs in the abdominal cavity (Johnson and Butterworth 1985). In adult flies, the fat body performs a variety of critical functions. As the primary metabolic organ, the fat body converts consumed nutrients into energy storage molecules like triglycerides and glycogen, and mobilizes these reserves as necessary to sustain the energetic needs of the animal (Canavoso et al. 2001; Arrese and Soulages 2010; Zheng et al. 2016). The fat body is also a highly secretory tissue, which releases a variety of molecules including proteins into the hemolymph. In female flies, the fat body produces vitellogenins, or “yolk” proteins which are secreted and taken up by the ovary to serve as a nutrient source for developing oocytes (Isaac and Bownes 1982; Minoo and Postlethwait 1985; Bownes and Blair 1986). In response to systemic microbial infections, the fat body is one of the main immunologically active tissues, expressing and secreting high levels of microbicidal immune effector molecules including antimicrobial peptides (AMPs; Tzou et al. 2000; Vaibhvi et al. 2022; Yu et al. 2022; Westlake et al. 2025). Because of this polyfunctionality and influence on other organ systems, the fat body plays an integral role in maintaining the overall health and physiological homeostasis of the animal. Moreover, the necessity of carrying out multiple, diverse biological processes simultaneously, often in response to dynamic extrinsic factors like dietary intake or infection, may cause the fat body to be particularly vulnerable to cellular strain.

The *yolk-GAL4* driver (also referenced as “*YP1-GAL4*”) is frequently used to conduct genetic manipulations in the adult fat body. In detail, the *P{yolk-GAL4}* transgene consists of a 1546 base pair sequence including the entire regulatory region between the vitellogenin-encoding genes *Yolk protein 1* (*Yp1*) and *Yolk protein 2* (*Yp2*) followed by *GAL4* coding sequence (Fig.1A; https://flybase.org/reports/FBsf0000933519; Georgel et al. 2001; Vidal et al. 2001). The native *Yp1* and *Yp2* genes are primarily expressed in the fat body and follicular epithelium of vitellogenic egg chambers in adult females (Gavin and Williamson 1976; Brennan et al. 1982; Isaac and Bownes 1982; Minoo and Postlethwait 1985; Bownes and Blair 1986). Accordingly, the *yolk-GAL4* transgene is only expressed in adult females (Fig.S1; Stronach et al. 2014), predominantly in the fat body and ovarian follicle cells (Fig.1B,B’,Fig.S1A-B’,I,I’). This expression pattern affords the technical advantage of inducing *UAS*-activated transgene expression only in the adult stage, thereby avoiding developmental lethality or other confounding effects that might result from expression of particular transgenes in the larval fat body. This inherent adult specificity of *yolk-GAL4* obviates the need for temporal control systems that rely on inducible de-repression or activation of GAL4 protein, such as *GAL80^TS^* (McGuire et al. 2003), *GAL80^AID^* (McClure et al. 2022), and GeneSwitch (Osterwalder et al. 2001), which all require temperature or dietary treatments that can have major impacts on fly physiology (Linder et al. 2008; Yamada et al. 2017; Klepsatel et al. 2019; Robles-Murguia et al. 2019; Ma et al. 2021; Gandara and Drummond-Barbosa 2022; Fleck et al. 2024). Despite these benefits and the extensive use of *yolk-GAL4* in over 70 published studies (https://flybase.org/reports/FBtp0014828.html), a few pieces of evidence have suggested potential deleterious effects from this transgene, particularly in relation to defense against pathogen challenge. Compared to non-transgenic flies, females carrying *yolk-GAL4* have been reported to display an attenuated innate immune response to bacterial infection (Kim et al. 2014) and to sustain higher viral titers when challenged with the insect-vectored Bluetongue virus (Shaw et al. 2012). However, the potential effects of *yolk-GAL4* on immunity or other fat body-regulated processes have not been carefully examined.

**Figure 1.**
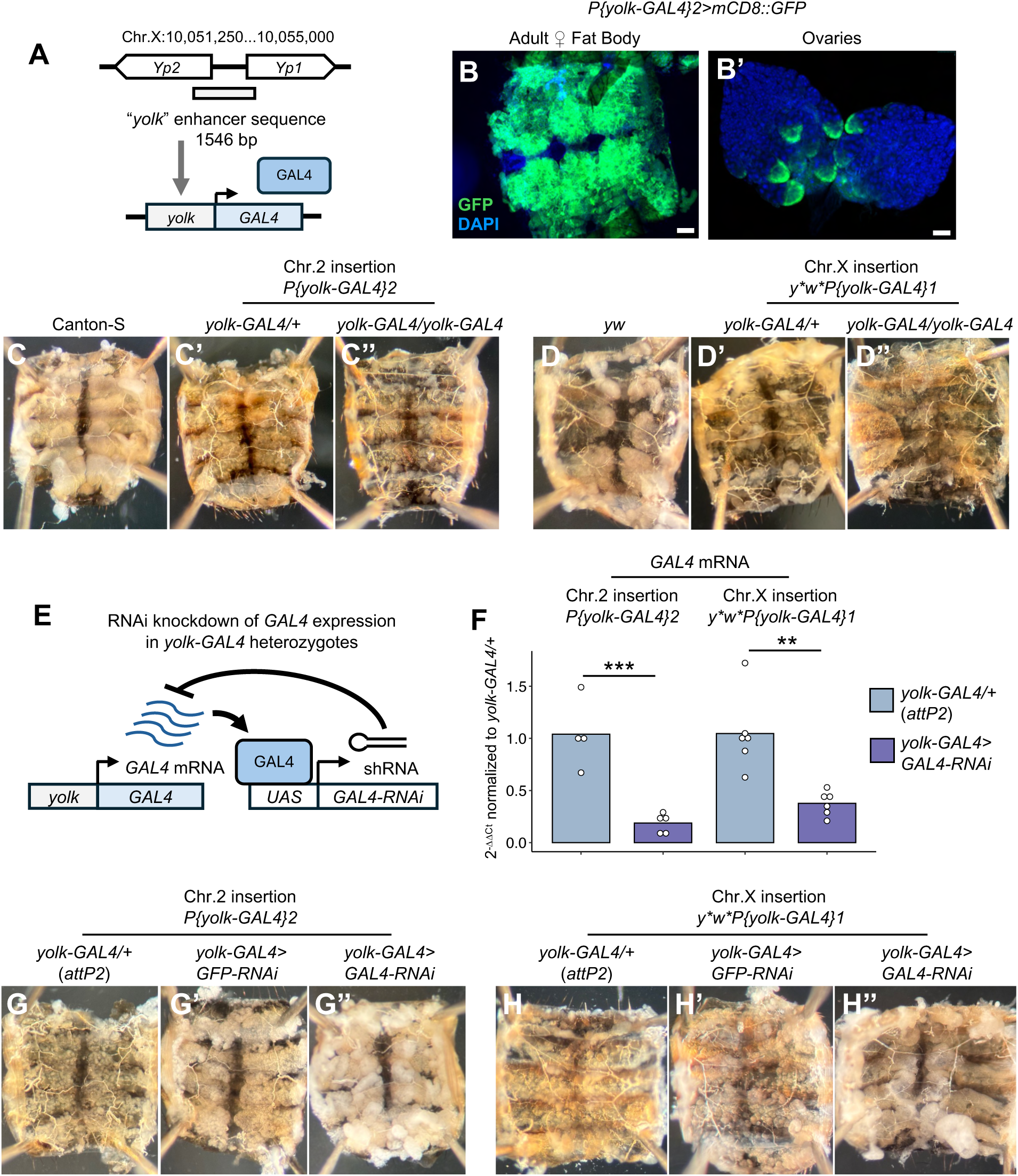
*GAL4* expression under control of the *yolk* regulatory sequence disrupts tissue integrity of the adult fat body. **(A)** Schematic illustrating design of the *yolk-GAL4* transgene (Vidal et al., 2001). Regulatory sequence spanning the intergenic region between the yolk protein-encoding genes *Yp1* and *Yp2* directs expression of the *S. cerevisiae* GAL4 transcription factor. **(B,B’)** Expression pattern of a *UAS-mCD8::GFP* reporter induced by *yolk-GAL4* indicates GAL4 expression in the adult female fat body **(B)** and follicle cells of maturing oocytes in the ovary **(B’)**. Nuclei are stained with DAPI to delineate overall tissue morphology. Scale bars=100µm. **(C-D’’)** Abdominal filets dissected from 6-7 day old adult females displaying fat body tissue lining the internal surface of the dorsal cuticle. Females heterozygous or homozygous for the same *yolk-GAL4* transgene integrated on either the second chromosome **(C’,C’’)** or the X chromosome **(D’,D’’)** contain thin, fragmented fat bodies, compared to robust adipose tissue in abdomens of non-transgenic Canton-S **(C)** and *yw* **(D)** control females. **(E)** Schematic illustrating genetic approach to reduce GAL4 expression levels in *yolk-GAL4* females. GAL4 protein produced from the *yolk-GAL4* transgene activates expression of a separate *UAS*-controlled transgene, which produces an RNA hairpin construct complementary to *GAL4* transcript sequence, thus leading to RNAi-mediated suppression of *GAL4* expression. **(F)** RT-qPCR analysis indicates that expressing the *UAS-GAL4-RNAi* construct substantially reduces *GAL4* expression levels in *yolk-GAL4* heterozygous females. Data represent 2^-11Ct^ values normalized to the mean ΔC_t_ value for control females with *yolk-GAL4* heterozygous over the *attP2* genetic background. Each dot represents an individual sample of 5-10 pooled flies, bars represent the mean. **p<0.01, ***p<0.001, unpaired two-sample t-test. **(G-H’’)** Females carrying *yolk-GAL4* heterozygous over the *attP*2 genetic background **(G, H)**, or expressing a *UAS-GFP-RNAi* construct (to control for non-specific effects of RNAi activation; **G’,H’**) contain fat bodies with morphological defects comparable to heterozygotes in **(C’,D’)**. Knocking down *GAL4* expression fully suppresses the aberrant fat body morphology caused by either the second chromosome **(G’’)** or X chromosome **(H’’)** *yolk-GAL4* insertion, indicating these phenotypes are caused by GAL4.

**Figure S1.**
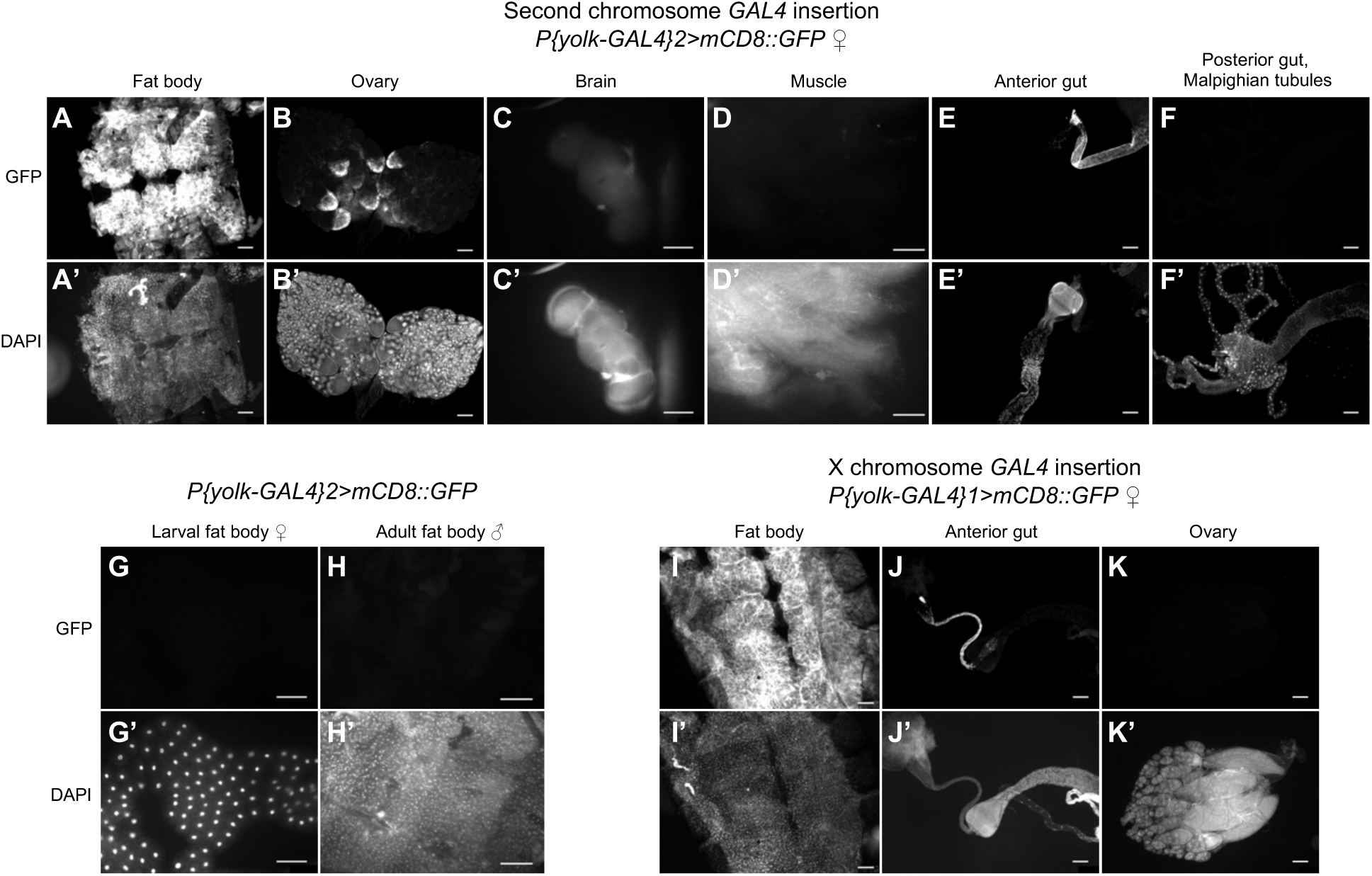
The *yolk-GAL4* transgene is expressed in the adult female fat body, ovarian follicle cells, and crop duct. Flies carrying the second chromosome *yolk-GAL4* insertion were crossed to a *UAS-mCD8::GFP* reporter line. Images represent tissues dissected from F1 progeny to assess *yolk-GAL4*-induced GFP expression. The second chromosome *yolk-GAL4* insertion activates strong GFP expression the fat tissue lining the abdominal cuticle **(A,A’)**, and in follicle cells surrounding maturing oocytes **(B,B’)**. Images in **(A-B’)** are the same as those presented in Fig.1B,B’, presented again here for direct comparison with other tissues and genotypes. **(C-F’)** No detectable live GFP signal is evident in the brain **(C,C’)**, muscle **(D,D’)**, anterior midgut **(E,E’)**, posterior midgut, or Malpighian tubules **(F,F’)**. Strong GFP expression is activated in the crop duct **(E,E’)**, which connects the esophagus to the crop. **(G-H’)** The *yolk-GAL4* transgene does not induce GFP expression in the fat body of larval females **(G,G’)**, or in adult male fat bodies **(H,H’)**, indicating the adult, female specificity of the driver. **(I-K’)** The *yolk-GAL4* transgene integrated on the X chromosome activates GFP expression in the adult female fat body **(I,I’)** and crop duct **(J,J’)**, but no GFP signal is detectable in the ovary (also see Fig.S3G-I). Nuclei are stained with DAPI to delineate overall tissue morphology. Scale bars=100µm.

Here we show that high-level *GAL4* expression activated by the *yolk* regulatory sequence severely compromises cellular and physiological functions of the adult *Drosophila* fat body. We find that flies carrying the *yolk-GAL4* transgene exhibit fat bodies with poor tissue integrity and reduced lipid stores, oogenesis defects with associated low fecundity, and a highly attenuated innate immune response associated with failure to survive bacterial infection. These phenotypes are all fully suppressed when we knock down *GAL4* expression by RNAi, indicating that GAL4 production directly causes these metabolic, reproductive, and immune defects. Comparing a panel of additional fat body driver lines, we find a strong, direct correlation between increasing fat body *GAL4* expression levels and greater likelihood of succumbing to bacterial infection. To determine whether the fat body defects we observed result from GAL4-specific toxicity or reflect more general effects of high levels of foreign protein in the fat body, we generated new *Drosophila* lines with the *yolk* regulatory sequence driving expression of cytoplasmic or nuclear-localized mCherry, or the alternative transactivator LexA. We find that only nuclear-localized mCherry and LexA proteins induce a transgene-dose-dependent reduction of infection survival, suggesting that excessive import or accumulation of transgenic proteins in fat body nuclei reduces females’ ability to withstand pathogen challenge.

## MATERIALS AND METHODS

### *Drosophila* stocks and husbandry

Genotypes and sources of all fly lines used in this study are detailed in Table S1. All stock and experimental flies were cultured on a diet consisting of (weight per volume or volume per volume) 6% cornmeal, 6% yeast, 4% sucrose, 0.7% agar, 0.265% methylparaben, 0.05% phosphoric acid, and 0.5% propionic acid. Flies were maintained at 25°C on a 12H:12H light:dark cycle. Except where indicated, all experiments were conducted with sibling-mated females aged 6-7 days post-eclosion.

Sequence data and information on fly stocks, transgenes, and gene function were obtained from FlyBase (release FB2025_02) (Jenkins et al. 2022).

### Tissue staining and fluorescence imaging

To visualize the expression patterns of fat body *GAL4* lines and *yolk-lexA* flies, males of the indicated driver genotype were crossed to virgin females carrying either *UAS-mCD8::GFP* or *LexAop-mCD8::GFP* to generate F1 progeny heterozygous for both the driver and responder elements. Expression patterns of the *yolk-mCherry.cyt* and *yolk-mCherry.nls* transgenes were visualized in tissues from flies homozygous for each transgene. Adult females were aged 6-7 days post-eclosion at 25°C (or 29°C for *3.1Lsp2>mCD8::GFP* flies; Fig.S4Ba-Bf) and dissected in ice-cold PBS (pH 7.4; Sigma) to isolate the brain, thoracic flight muscle, ovaries, gut, Malpighian tubules, and abdominal carcass. Larval fat bodies were dissected at the late third instar stage, ∼96 hours post egg-laying. Tissues were fixed in 4% paraformaldehyde (Electron Microscopy Sciences) for 30 min, washed three times in PBS 0.3% Triton X-100 (PBT), and stained with DAPI (1µg/mL in PBS; Invitrogen) for 15 min at room temperature to label cell nuclei. Tissues were washed in PBS and mounted in ProLong Glass Antifade Mountant (Invitrogen).

Epifluorescence images were acquired on a Leica DM5000B upright microscope with a CTR5000 controller, EL6000 light source, and DFC365 FX camera using LAS AF 2.6 software.

### Oil Red O staining and image analysis

Abdominal fat tissue was visualized by Oil Red O staining following established protocols (Gutierrez et al. 2007; Molaei et al. 2019). Dissected abdominal carcasses were fixed as described above and stained for 30min in freshly prepared Oil Red O solution: 6mL 0.1% Oil Red O (Sigma) dissolved in isopropanol and 4mL distilled water passed through a 0.22μm filter. Carcasses were rinsed three times with distilled water, mounted as described above, and imaged on a Leica M165 FluoCombi stereomicroscope system.

Oil Red O staining intensities were quantified using FIJI. Color images were converted to 8-bit pixel depth and inverted. For each carcass, a constant 477×525 pixel region of interest was positioned over abdominal segments A2-A5 (Yoder 2012; Jürgens et al. 2024) and mean pixel intensity was measured within this area.

### Starvation survival analysis

Flies were transferred to 1% (weight per volume) agarose-water vials and maintained at 25°C on a 12H:12H light:dark cycle. Mortality was scored three times per day until the entire population succumbed.

### Egg laying and hatch rate assays

For egg laying assays, females were collected as virgins 0-4 hours post-eclosion and placed in food vials in groups of two or three flies per vial with an equal number of Canton-S males. Flies were flipped to new food vials every 24 hours. Total eggs laid in each 24 hour time period were counted and divided by the number of females in the vial to approximate the number of eggs laid per fly. These daily values were summed to calculate total eggs laid per fly over the 7 day experimental period.

To assay embryo viability, the numbers of eggs laid in food vials over a 24 hour period were counted and resulting progeny were allowed to develop to the pupal stage for 8 days 25°C. Hatch rates were calculated as the ratio of the number of pupae to the number of eggs laid in each vial.

### Ovary size and egg chamber stage analyses

Whole intact ovaries were dissected in PBS and immediately imaged on a Leica M165 FluoCombi stereomicroscope system. Ovary size was quantified in FIJI, using the polygon selection tool to trace the outline of each ovary image and measure the area within the outlined region.

Numbers of vitellogenic follicles and mature oocytes per ovary were counted by visual inspection of dissected ovaries under a stereomicroscope. Vitellogenic follicles (stages 8-13) were identified by the presence of opaque, yolk-filled oocytes morphologically distinguishable from nurse cells (Jia et al. 2016). Mature, stage 14 oocytes were identified by the presence of fully-formed dorsal appendages (Bastock and St Johnston 2008).

### Bacterial infection procedures and infection survival analysis

Flies were systemically infected with *Providencia rettgeri* strain Dmel (Juneja and Lazzaro 2009), *Enterococcus faecalis* Dmel (Lazzaro 2002), or *Pectobacterium carotovorum* (Ecc15; Basset et al. 2000) following procedures described in Khalil et al. (2015). Flies were briefly anesthetized on CO_2_ platforms and *P. rettgeri* and *E. faecalis* infections were administered by dipping a 0.15mm minutien pin in the bacterial inoculum and pricking flies in the sternopleural region of the thorax. *Ecc15* infections were administered by injecting 23nL of bacterial suspension into the thorax with a Drummond Nanoject II nanoinjector. Bacterial cultures were grown overnight in either Luria-Bertani (LB) broth (*P. rettgeri* and *Ecc15*) or Brain Heart Infusion (BHI) broth (*E. faecalis*). *P. rettgeri* cultures were grown at 37°C with shaking, and *E. faecalis* and *Ecc15* cultures were grown at 30°C with shaking. Overnight cultures were pelleted by centrifugation and resuspended in sterile phosphate-buffered saline (PBS). *P. rettgeri* and *E. faecalis* inocula were standardized to optical density (OD_600_)=1, corresponding to ∼300 colony forming units (CFU) per pinprick, while *Ecc15* was standardized to OD_600_=10, corresponding to ∼10^5^ CFU injected into each fly. To quantify bacterial doses administered with each infection, 4-8 flies were infected and immediately homogenized and plated to enumerate CFUs as described below.

Infected flies were housed in food vials in groups of 10-15 individuals per vial at 25°C and flipped to fresh vials every other day due to growth of larval progeny in the food substrate. To quantify infection-induced mortality, the number of dead flies in each vial was counted daily for five days post-infection.

### Bacterial load analysis

Individual flies were collected at the indicated timepoints post-infection and homogenized in 250μL PBS with 2.3mm chrome steel disruption beads (Research Products International) via bead-beating for 30 seconds with a Mini-BeadBeater-96 tissue disrupter (BioSpec Products). Homogenates were diluted 1:10 and 1:100 in PBS, and each dilution was plated on LB agar with a Whitley Automated Spiral Plater 2 (Don Whitley Scientific). Plates were incubated at 37°C overnight and resulting colonies were counted from dilution plates that yielded distinct, individual colonies using a ProtoCOL3 plate counter (Microbiology International). CFU per ml values obtained from the ProtoCOL3 software were used to calculate the number of viable bacterial cells in each fly with the following equation: CFU/fly = (CFU/mL) * (dilution factor) * (0.25mL)

### RT-qPCR and RT-PCR

Whole flies or dissected tissues were homogenized for 30 seconds in 500μL Trizol reagent (Invitrogen) with 2.3mm zirconia/silica beads (BioSpec Products) via bead-beating. Total RNA was extracted using the Zymo Research Direct-zol RNA miniprep kit and cDNA was synthesized from 500ng RNA template with Bio-Rad iScript Reverse Transcription Supermix. qPCR was performed with PerfeCTa SYBR Green FastMix (Quantabio) on a Bio-Rad CFX Connect thermocycler. Expression levels for genes of interest were calculated as ΔΔCt values:

_1)_ For each sample, Ct values of the housekeeping gene *Rpl32* were subtracted from Ct values of the gene of interest: ΔCt_GOI_=Ct_GOI_ – Ct_Rpl32_
2) Experimental genotypes/conditions were normalized to control genotypes/conditions by first averaging the ΔCt values for the control group and then subtracting this average value from the ΔCt values of each individual experimental sample: ΔΔCt_GOI_=ΔCt_experimental_sample_ – average(ΔCt_control_samples_)

For RT-PCR confirmation of *UAS-p35* transgene expression (Fig.S6A), RNA was extracted and cDNA synthesized as described above. PCR amplification was conducted from cDNA template using GoTaq Green Master Mix (Promega) and the following cycling conditions: 2 min initial denaturation at 95°C, followed by 30 cycles of 30 sec denaturation at 95°C, 30 sec annealing at 50°C, and 1 min elongation at 72°C. Reaction products were resolved on a 1.5% agarose gel with SYBR Safe DNA gel stain (Invitrogen).

Primer sequences are listed in Table S2.

### Generation of yolk-mCherry.cyt and yolk-mCherry.nls lines

*pYolk-mCherry.NLS-attB* and *pYolk-mCherry.CYT-attB* plasmids were generated by VectorBuilder Inc. The *yolk* regulatory sequence used for construction of the *yolk-GAL4* transgene (https://flybase.org/reports/FBsf0000933519) was cloned upstream of *mCherry* coding sequence, replacing the *UASt-Hsp70p* regulatory sequence in the *pUASt.attB* vector backbone. In the *pYolk-mCherry.NLS-attB* construct, *mCherry* is flanked by 5’ and 3’ nuclear localization signal sequences. Constructs were injected into *M{vas-int.Dm}ZH-2A;P{CaryP}attP40* embryos by BestGene, Inc. for ΦC31-mediated transgene integration at the second chromosome *attP40* docking site. Positive transformants were selected by expression of a *mini-white* eye marker, balanced, and established as stable lines in a *w^1118^* background.

### Generation of *yolk-LexA* by HACKy-mediated conversion of *yolk-GAL4*

The second chromosome *yolk-GAL4* insertion was converted to *yolk-LexA* following the ‘version 2’ HACKy approach detailed in Rankin et al. (2024). The crossing scheme employed for conversion is presented in Fig.S9A. First, *yolk-GAL4* males (expressing a *mini-white*^+^ visible marker) were crossed to females carrying germline-expressed *vas-Cas9* transgene and the HACKy donor cassette (abbreviated “*CyO, PBac{LexA.G4H, RFP^+^, y^+^}*”). The HACKy donor comprises: 1) U6 promoter-expressed guide RNAs targeting the *GAL4* coding sequence, 2) the *LexA::GAD* coding sequence flanked by *GAL4* homology arms and separated from the 5’ homology arm by T2A ribosomal skipping sequence, and 3) the *3xP3-RFP* and *y^+t7.7^* visible markers. Eighty individual F1 males, carrying *vas-Cas9* on the X chromosome and *yolk-GAL4* in trans to *CyO, PBac{LexA.G4H, RFP^+^, y^+^}* were mated to 2-3 *y^1^w^67c23^;sna^ScO^/CyO* virgin females each. We screened ∼50-100 F2 progeny from each F1 cross and identified a single, balanced male expressing all three *mini-white*^+^, *y^+^*, and *3xP3-RFP* markers, indicative of successful conversion. This male was sequentially crossed to *y^1^w^67c23^;sna^ScO^/CyO* virgin females to establish a stable line, and then to *LexAop-mCD8::GFP* virgin females to confirm GFP expression in the spatiotemporal pattern matching that activated by *yolk-GAL4*.

### Statistical analyses

All statistical tests were performed with R version 4.4.0 (https://R-project.org). For all experiments except survival analyses, data normality was first evaluated by Shapiro-Wilk test and homogeneity of variance assessed by Levene’s test. Data that met parametric test assumptions (normal distribution and equal variance among groups) were analyzed by unpaired, two-sample t-test (two experimental genotypes), or by one-way ANOVA with Tukey’s post hoc comparisons (three or more genotypes). Data that did not meet parametric test assumptions were compared via Mann-Whitney test or Kruskal-Wallis test with Dunn’s post hoc comparison. Figure legends indicate the test used for each experiment. Infection and starvation survival data were analyzed via pairwise Log-Rank test with Bejamini-Hochberg correction using the pairwise_survdiff() function in the survminer package (Kassambara et al. 2024). The significance threshold was considered p<0.05. Sample sizes for each dataset are reported in Table S3.

## RESULTS

### GAL4 expression under control of the *yolk* regulatory sequence disrupts tissue morphology of the adult female fat body

The *yolk-GAL4* transgene directs GAL4 expression specifically in the adult female fat body (Fig.1B, Fig.S1A,A’,I,I’) and in the follicle cells of late-stage oocytes (Fig.1B’, Fig.S1B,B’) with additional activity observed in the crop duct (Fig.S1E,E’,J,J’). While attempting to conduct genetic manipulations with a fly line carrying *yolk-GAL4* integrated on the second chromosome (*P{yolk-GAL4}2*), we noticed that the adipose tissue of flies harboring this construct exhibited striking morphological defects. Fat bodies lining the dorsal abdomen of *P{yolk-GAL4}2* heterozygotes appeared thin, fragmented, and clumpy (Fig.1C’) compared to the robust sheets of adipose tissue in wild-type Canton-S flies (Fig.1C). We observed comparable aberrant tissue integrity in fat bodies of *P{yolk-GAL4}2* homozygous females (Fig.1C’’).

These observations suggested that *GAL4* expressed under control of the *yolk* promoter could disrupt fat body tissue integrity. However, the *P{yolk-GAL4}2* line was generated by P-element-mediated integration at an unmapped genetic locus on the second chromosome (Georgel et al. 2001; Vidal et al. 2001). Thus, in principle, the *yolk-GAL4* insertion might disrupt expression or function of a gene necessary to maintain fat body tissue integrity. To address this possibility, we examined a separate fly line carrying the same *P{yolk-GAL4}* construct integrated on the X chromosome (*P{yolk-GAL4}1*), and therefore at an entirely different genomic locus. Females carrying one or two copies of *P{yolk-GAL4}1* contained fragmented fat bodies morphologically similar to those observed in *P{yolk-GAL4}2* females (Fig.1D’,D’’). These results strongly suggest that the aberrant fat body morphology of *yolk-GAL4* flies is caused by expression of the transgene, and not by inadvertent insertional mutagenesis of an unidentified gene.

We reasoned that if the aberrant fat body morphology we observed in *yolk-GAL4* females were truly caused by GAL4, then blocking *GAL4* expression should alleviate this defect. To test this prediction, we generated *yolk-GAL4* heterozygotes with self-targeted, RNAi-mediated inhibition of *GAL4* expression (Fig.1E). Specifically, we crossed each of the two *yolk-GAL4* insertion lines to flies carrying a *UAS*-activated RNA hairpin construct targeting the *GAL4* coding sequence. Expression of GAL4 protein therefore results in RNAi knockdown of the *GAL4* transgene in F1 progeny (Fig.1E). For genetic controls, we crossed *yolk-GAL4* flies with the genetic background line in which the *UAS-GAL4-RNAi* stock was generated (“*attP2*”), and with flies carrying a *UAS-GFP-RNAi* construct to control for non-specific effects of RNAi activation. RT-qPCR analysis confirmed that *GAL4* mRNA levels were substantially reduced when *GAL4* RNAi was activated by either *P{yolk-GAL4}2* or *P{yolk-GAL4}1* (Fig.1F).

We found that *GAL4* knockdown fully suppressed the aberrant fat body morphology caused by heterozygosity for *yolk-GAL4*. Females carrying either *yolk-GAL4* insertion heterozygous over the *attP2* background, or with *yolk-GAL4* activating expression of the *UAS-GFP-RNAi* construct contained thin, clumpy fat bodies similar to those observed with *yolk-GAL4* in *trans* to Canton-S or *yw* backgrounds (Fig.1G,G’,H,H’). By contrast, females heterozygous for *yolk-GAL4* with RNAi-mediated *GAL4* knockdown displayed robust, contiguous fat body sheets comparable to those observed in flies lacking the transgene (Fig.1G’’,H’’). Taken together, these results indicate that *GAL4* expression activated by the *yolk* regulatory sequence compromises the gross-level tissue integrity of the adult fat body.

### *yolk-GAL4* expression impairs major physiological functions of the adult fat body

Having found that *yolk-GAL4* compromises adipose tissue integrity, we hypothesized that expression of this transgene might also impact major physiological processes regulated by the adult female fat body, namely energy storage, immunity, and reproduction.

#### *yolk-GAL4* expression reduces lipid stores and starvation resistance

Females heterozygous or homozygous for *yolk-GAL4* contained thin, fragmented fat body tissue (Fig.1C’,C’’,D’,D’’,G,G’,H,H’). We therefore asked whether this degenerated tissue appearance might be accompanied by reduced levels of stored lipids. To answer this question, we visualized gross fat levels in wild-type, *yolk-GAL4/+* and *yolk-GAL4/yolk-GAL4* females by staining dissected abdominal carcasses with the lipophilic dye Oil Red O (ORO). ORO non-specifically labels neutral lipids including triglycerides (Ramírez-Zacarías et al. 1992), which constitute the primary class of lipids in the *Drosophila* fat body (Canavoso, Jouni, K.J. Karnas, et al. 2001; Parisi et al. 2011; Kühnlein 2012). We found that one or two copies of either *P{yolk-GAL4}2* (Fig.2A’,A’’) or *P{yolk-GAL4}1* (Fig.S2A’,A’’) resulted in significantly weaker ORO staining intensity compared to Canton-S (Fig.2A) or *yw* (Fig.S2A) controls (Fig.2B, Fig.S2B). Reduced fat levels are frequently accompanied by greater sensitivity to full nutrient deprivation (Da Lage et al. 1989; Zwaan et al. 1991; Chippindale et al. 1996; Gibbs and Reynolds 2012; Chauhan et al. 2021). Consistent with their reduced abdominal lipid content, we found that females carrying *yolk-GAL4* succumbed to starvation considerably more rapidly than non-transgenic controls (Fig.2C, Fig.S2C). Notably, for both *yolk-GAL4* insertion lines, heterozygotes survived moderately but significantly longer than homozygotes (Fig.2C, Fig.S2C), suggesting a dose-dependent effect of *GAL4* on starvation sensitivity.

**Figure 2.**
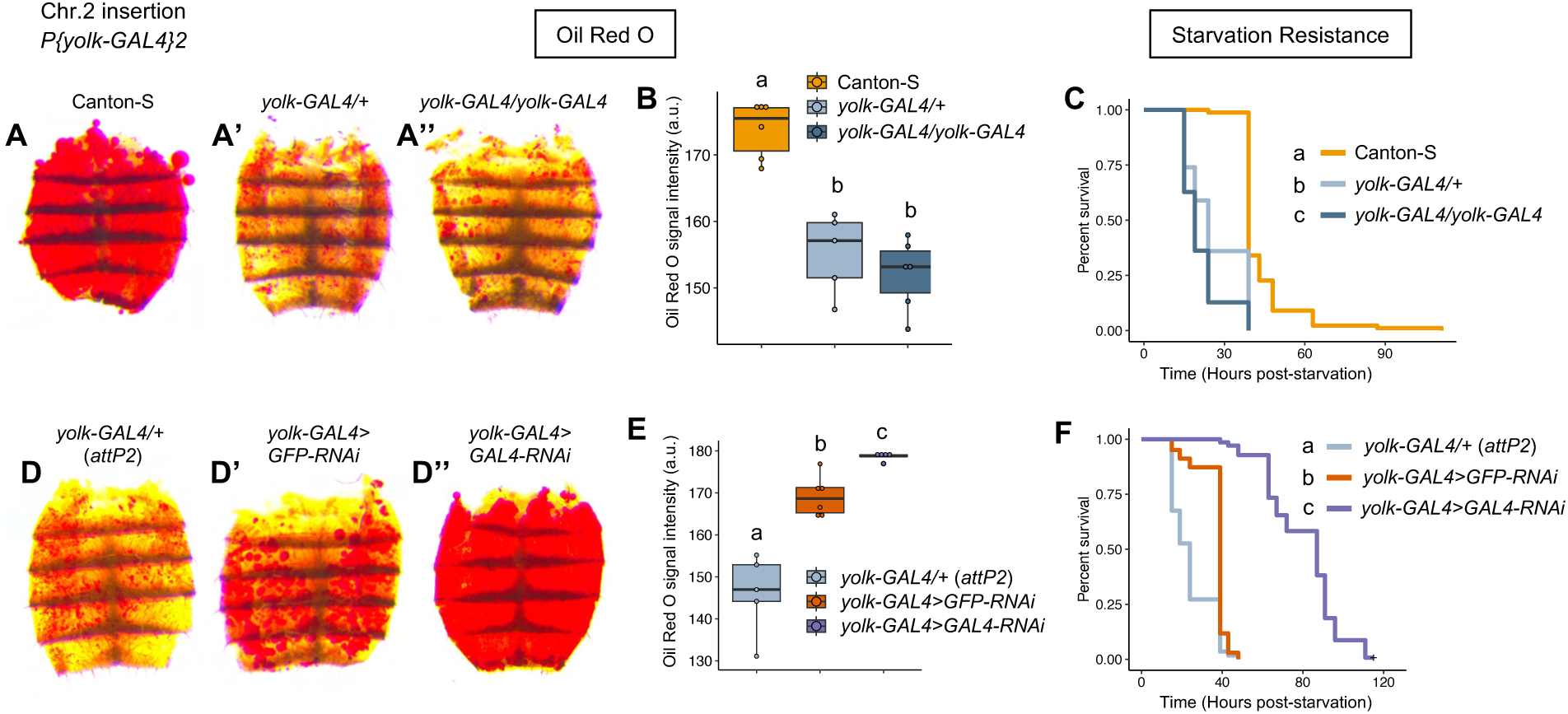
Expression of *yolk-GAL4* reduces fat body lipid stores and increases starvation sensitivity. (A-A’’) Representative images of abdominal filets dissected from 6-7 day old adult females of the indicated genotypes, stained with the lipophilic dye Oil Red O (ORO). Females heterozygous **(A’)** or homozygous **(A’’)** for the second chromosome *yolk-GAL4* insertion display weaker abdominal ORO staining compared to non-transgenic Canton-S controls **(A)**. **(B)** Quantification and statistical comparisons of ORO staining intensities. **(C)** Females carrying *yolk-GAL4* succumb to starvation faster than non-transgenic control females. **(D-F)** Knocking down *GAL4* expression by using *yolk-GAL4* to induce *GAL4*-targeting RNAi (see Fig.1E,F) increases abdominal fat levels **(D-D’’,E)** and extends starvation survival time **(F)** in females carrying *yolk-GAL4*. For ORO signal intensity data **(B,E)** each dot represents an individual abdomen/fly, a.u.=arbitrary units. Statistics: ORO data were analyzed by one-way ANOVA with Tukey’s post-hoc pairwise comparisons **(B,E)**. Starvation survival data were analyzed by pairwise Log-Rank tests with Benjamini Hochberg correction for multiple comparisons **(C,F)**. Genotypes with different letters are statistically different from one another (p<0.05). All sample sizes are reported in Table S3.

**Figure S2.**
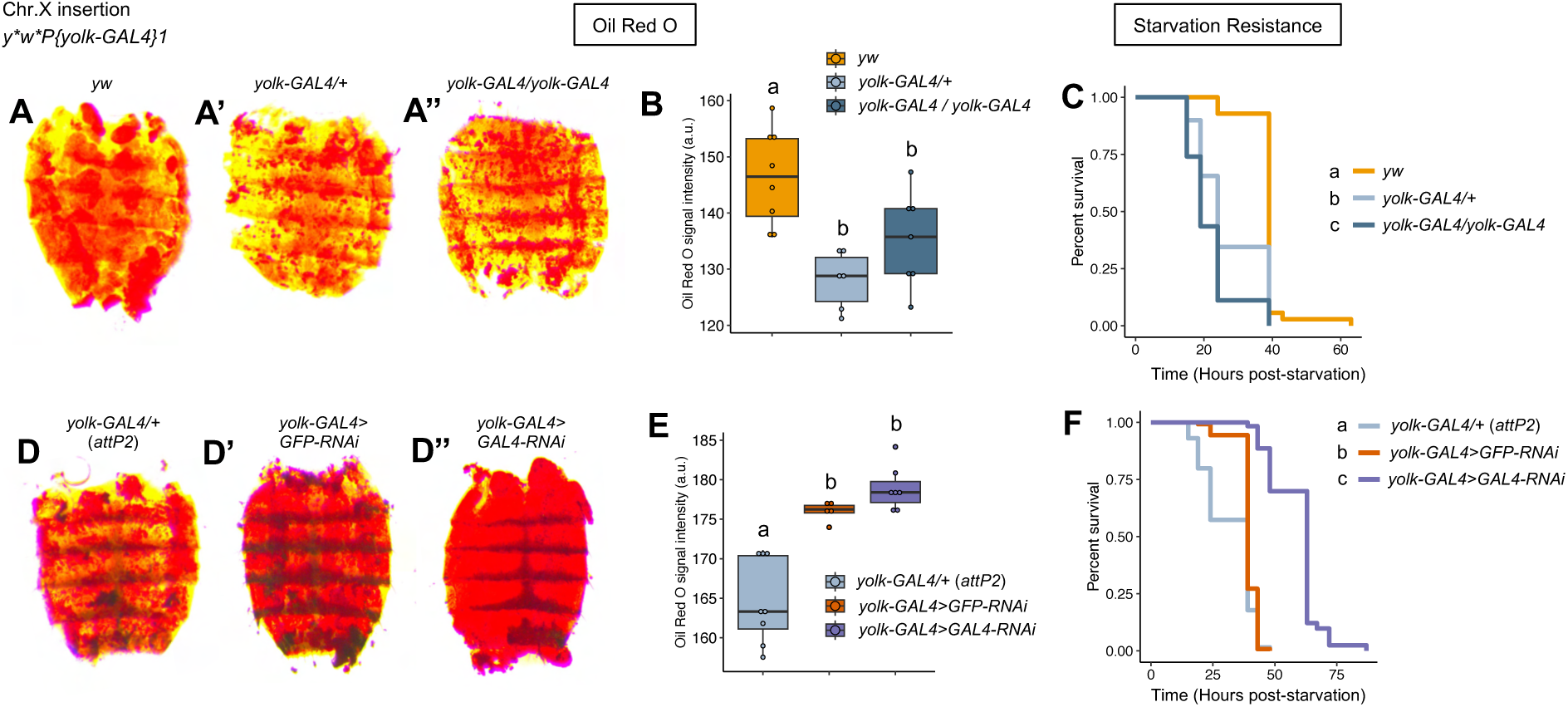
The *yolk-GAL4* transgene integrated on the X chromosome decreases fat stores and starvation resistance. (A-A’’) Representative images of ORO-stained abdominal filets dissected from 6-7 day old adult females of the indicated genotypes. Compared to non-transgenic *yw* background controls **(A)**, females carrying one **(A’)** or two copies **(A’’)** of the X chromosome *yolk-GAL4* insertion display weaker abdominal ORO staining. **(B)** Quantification and statistical comparisons of ORO staining intensities. **(C)** Heterozygosity or homozygosity for *yolk-GAL4* on the X chromosome increases starvation sensitivity. **(D-F)** Reducing *GAL4* expression by RNAi increases abdominal lipid levels **(D-E)** and starvation resistance **(F)** of flies with X chromosome *yolk-GAL4*, similarly to effects of knocking down *GAL4* expressed from the second chromosome insertion (see Fig.2D-F). For ORO signal intensity data **(B,E)** each dot represents an individual abdomen/fly, a.u.=arbitrary units. Statistics: ORO data were analyzed by one-way ANOVA with Tukey’s post hoc pairwise comparisons **(B)** or by Kruskal-Wallis test with Dunn’s post hoc pairwise comparisons **(E)**. Starvation survival data were analyzed by pairwise Log-Rank tests with Benjamini Hochberg correction for multiple comparisons **(C,F)**. Genotypes with different letters are statistically different from one another (p<0.05).

To confirm that GAL4 caused the low fat content and starvation sensitivity of *yolk-GAL4* flies, we again reduced *GAL4* expression levels by RNAi and quantified ORO staining and starvation resistance. We found that knocking down *GAL4* expression from either *P{yolk-GAL4}2* or *P{yolk-GAL4}1* led to higher fat levels (Fig.2D’’, Fig.S2D’’) and increased starvation resistance (Fig.2F, Fig.S2F) compared to control animals with *yolk-GAL4* heterozygous over the *attP2* background (Fig.2D,E Fig.S2D,E) or with *yolk-GAL4* activating expression of the *UAS-GFP-RNAi* construct (Fig.2D’,E, Fig.S2D’,E). Of note, for both insertions, *yolk-GAL4>GFP-RNAi* females displayed less severe fat depletion and starvation sensitivity than *yolk-GAL4/attP2* females (Fig.2D,D’,E,F, Fig.S2D,D’,E,F), suggesting the *UAS-GFP-RNAi* genetic background may harbor elements capable of partially suppressing these phenotypes.

Collectively these data show that expression of the *yolk-GAL4* transgene results in diminished abdominal fat stores and increased sensitivity to starvation, both indicative of disrupted metabolic homeostasis.

#### *yolk-GAL4* impairs oogenesis and egg laying

*yolk-GAL4* is expressed in the fat body and ovarian follicle cells (Fig.1B,B’, Fig.S1A,B,I), both of which regulate oogenesis (Brennan et al. 1982; Isaac and Bownes 1982; Arrese and Soulages 2010). We therefore asked whether *yolk-GAL4* might affect egg production. To answer this question, we counted the total numbers of eggs laid by control females and females carrying *P{yolk-GAL4}2* for seven days post-eclosion. This analysis revealed major egg laying deficits caused by the *P{yolk-GAL4}2* transgene. Over the first week of adulthood, *P{yolk-GAL4}2* heterozygous or homozygous females laid on average fewer than 50 eggs, in contrast to the ∼200 eggs laid by Canton-S wild-type flies (Fig.3A). Reduced egg laying could result from decreased oogenesis or from retention of mature oocytes and reduced ovulation. Visual inspection of ovaries dissected from *P{yolk-GAL4}2* heterozygous (Fig.3B,C’) and homozygous (Fig.3B,C’’) females revealed them to be dramatically stunted in size compared to the fully developed ovaries of Canton-S flies (Fig.3B,C), suggesting that the low egg output of *P{yolk-GAL4}2* females likely reflects oogenesis defects as opposed to reduced ovulation.

**Figure 3.**
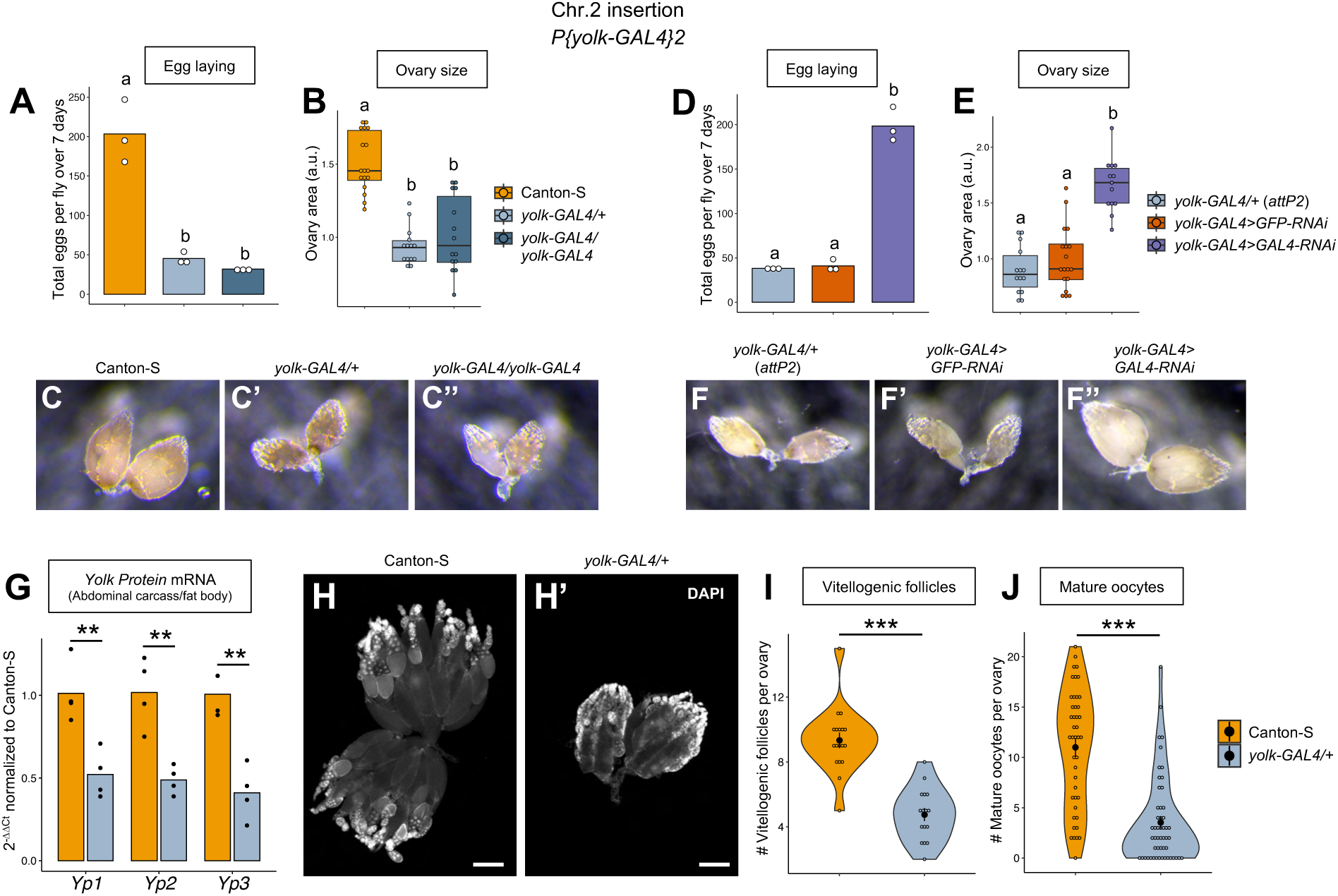
Females expressing *yolk-GAL4* exhibit oogenesis and egg laying defects. **(A)** Females heterozygous or homozygous for *yolk-GAL4* lay dramatically fewer eggs compared to non-transgenic Canton-S females. **(B-C’’)** Ovaries from females carrying one or two copies of *yolk-GAL4* are stunted in size compared to ovaries from wild-type, Canton-S females. Representative images of whole dissected ovaries are presented in **(C-C’’)**, a.u.=arbitrary units. **(D-F’’)** Reducing *GAL4* expression by RNAi fully suppresses the egg laying deficit **(D)** and ovary stunting **(E-F’’)** caused by heterozygosity for *yolk-GAL4*. For egg laying data **(A,D)**, each point represents an individual replicate, 2-3 females per replicate, bars represent the mean. For quantification of ovary size **(B,E)**, each point represents an individual ovary. Box plots represent the median, first and third quartiles, and 1.5x interquartile range. **(G)** RT-qPCR analysis shows that expression of the three major Yolk Protein-encoding genes is reduced in fat bodies of females heterozygous for *yolk-GAL4*. Data represent 2^-11Ct^ values normalized to the mean ΔC_t_ value for Canton-S samples. Each dot represents an individual sample of 10 pooled fat bodies/abdominal carcasses, bars represent the mean. **(H,H’)** Ovaries dissected from non-transgenic Canton-S females **(H)** and *yolk-GAL4* heterozygous females **(H’)** with DAPI-stained nuclei to reveal developmental stages of individual egg chambers. Scale bars=100µm. **(I-J)** Ovaries from *yolk-GAL4* heterozygotes contain fewer vitellogenic egg chambers **(I)** and fewer mature oocytes **(J)** than ovaries from wild-type females, suggesting *yolk-GAL4* expression impairs yolk deposition and egg maturation. For both violin plots, each point represents counts from an individual ovary. Opaque point represents the mean and bars represent standard error. Statistics: Egg laying data **(A,D)** were analyzed by one way ANOVA with Tukey’s post hoc pairwise comparisons. Ovary size data were analyzed by Kruskal-Wallis test with Dunn’s post hoc pairwise comparisons **(B)** or by one way ANOVA with Tukey’s post hoc pairwise comparisons **(E)**. Genotypes with different letters are statistically different from one another (p<0.05). RT-qPCR data **(G)** and vitellogenic egg chamber counts **(I)** were analyzed by unpaired two-sample t-test. Mature oocyte counts **(J)** were analyzed by Mann-Whitney test. **p<0.01, ***p<0.001.

**Figure S3.**
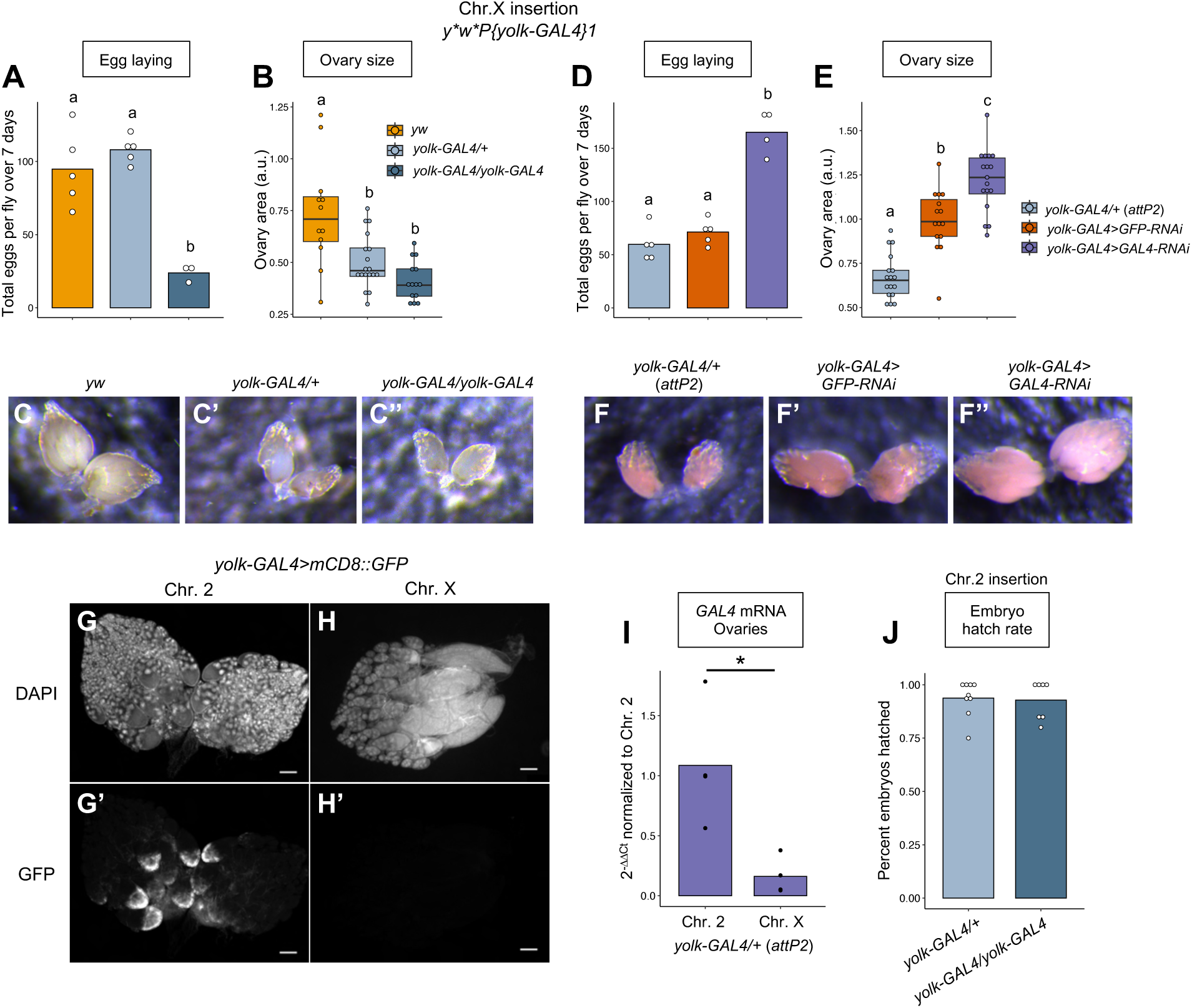
The X chromosome *yolk-GAL4* insertion is minimally expressed in the ovary and induces weaker oogenesis defects. **(A)** Females heterozygous for the *yolk-GAL4* insertion on the X chromosome lay comparable numbers of eggs to non-transgenic *yw* controls, while *yolk-GAL4* homozygotes have a lower egg output. **(B-C’’)** Ovaries of females heterozygous or homozygous for *yolk-GAL4* on the X chromosome are reduced in size compared to ovaries of *yw* females. **(D-F’’)** RNAi knockdown of *GAL4* expressed from the X chromosome *yolk-GAL4* insertion increases egg output **(D)** and ovary size **(E-F’’)**. For egg laying data **(A,D)**, each point represents an individual replicate, 2-3 females per replicate, bars represent the mean. For quantification of ovary size **(B,E)**, each point represents an individual ovary. **(G-H’)** The *yolk-GAL4* transgene integrated on the second chromosome activates *UAS-mCD8::GFP* reporter expression in the follicle cells **(G,G’)**, whereas the same transgene integrated on the X chromosome does not produce visible GFP signal **(H,H’)**. These are the same images as in Fig.1B’ and Fig.S1B,B’,K,K’ presented again here for direct comparison between genotypes. Scale bars=100µm. **(I)** RT-qPCR analysis reveals reduced *GAL4* transcript levels in ovaries of females heterozygous for the X chromosome *yolk-GAL4* insertion compared to the second chromosome *yolk-GAL4* insertion. Data represent 2^-11Ct^ values normalized to the mean ΔC_t_ value for Canton-S samples. Each dot represents an individual sample of 10 pooled ovaries, bars represent the mean. **(J)** The majority of eggs laid by *yolk-GAL4* heterozygous or homozygous females successfully hatch, indicating no major quality problems despite the low numbers of eggs produced. Statistics: Egg laying data **(A,D)** were analyzed by one way ANOVA with Tukey’s post hoc pairwise comparisons. Ovary size data were analyzed by Kruskal-Wallis test with Dunn’s post hoc pairwise comparisons **(B)** or by one way ANOVA with Tukey’s post hoc pairwise comparisons **(E)**. RT-qPCR data **(I)** were analyzed by unpaired, two-sample t-test. *p<0.05.

To test whether *GAL4* expression from the *yolk* enhancer leads to reduced egg laying and stunted ovaries, we again examined the X chromosome *P{yolk-GAL4}1* line and knocked down *GAL4* expression by RNAi. While *P{yolk-GAL4}1* homozygotes laid significantly fewer eggs than non-transgenic controls, heterozygosity for *P{yolk-GAL4}1* did not affect egg output (Fig.S3A). Of note, however, *yw* females used as the control genotype for this experiment laid substantially fewer eggs than the Canton-S females used as controls for *P{yolk-GAL4}2* (compare orange bars, Fig.3A and Fig.S3A). Despite these different egg laying rates, both *P{yolk-GAL4}1* heterozygotes and homozygotes contained significantly smaller ovaries than *yw* flies (Fig.S3B-C’’). Moreover, we found that knocking down *GAL4* levels fully suppressed the low egg output and reduced ovary size of females heterozygous for *P{yolk-GAL4}2* (Fig.3D-F’’) or *P{yolk-GAL4}1* (Fig.S3D-F’’), confirming that expression of the *yolk-GAL4* transgene leads to decreased egg production.

Across multiple genotypes, we noticed that females heterozygous for *P{yolk-GAL4}2* laid fewer eggs than those heterozygous for *P{yolk-GAL4}1* (compare corresponding colors between Fig.3A,D and Fig.S3A,D). We speculated that this difference in phenotype severity could reflect differences in GAL4 expression strength or pattern resulting from the distinct chromosomal insertion sites of the transgene between these two lines. Consistent with this, while both insertions drove robust expression of a *UAS-GFP* reporter in the fat body and crop duct (Fig.S1A,E,I,J), we found that only *P{yolk-GAL4}2* activated visible GFP expression in the ovary (Fig.S1B,B’,K,K’, Fig.S3G-H’). RT-qPCR analysis confirmed that *GAL4* expression was significantly lower in ovaries of *P{yolk-GAL4}1* heterozygotes compared to *P{yolk-GAL4}2* heterozygotes (Fig.S3I). These data suggest that the quantitative level *GAL4* expression in the follicular epithelium might contribute to the severity of egg laying defects caused by *P{yolk-GAL4}2*.

We next asked whether ovary stunting and reduced egg laying induced by *GAL4* expression might reflect vitellogenesis defects originating from impaired yolk protein synthesis by the fat body. To test this hypothesis, we used RT-qPCR to measure *Yp1*, *Yp2*, and *Yp3* expression levels in fat bodies dissected from Canton-S and *P{yolk-GAL4}2*/+ females. We found that expression of all three *yolk* genes was significantly reduced in females carrying *P{yolk-GAL4}2* (Fig.3G), suggesting the transgene might curtail the fat body’s ability to provide vitellogenins to developing oocytes. Closer inspection of dissected ovaries revealed that ovarioles from *P{yolk-GAL4}2/+* females contained substantially fewer vitellogenic follicles (Fig.3H,H’,I) and fewer mature oocytes (Fig.3H,H’,J), suggesting impaired yolk provisioning and consequent oogenesis failure. Notably, however, >94% of the eggs laid by *P{yolk-GAL4}2* heterozygous or homozygous females yielded viable progeny, suggesting no major decline in the quality of the few oocytes that are successfully produced (Fig.S3J).

Altogether, these findings reveal that expression of the *yolk-GAL4* transgene interferes with vitellogenesis, leading to substantially curtailed egg production.

#### *yolk-GAL4* compromises resistance to systemic bacterial infections

A previous study by Kim et al. (2014) reported abrogated transcriptional upregulation of genes encoding antimicrobial peptides following *E. coli* infection in *yolk-GAL4* heterozygotes compared to non-transgenic *w^1118^* flies. Based on this prior study, and because the fat body is the principal tissue responsible for AMP production during systemic microbial infections (Tzou et al. 2000; Arrese and Soulages 2010; Westlake et al. 2025), we hypothesized that *yolk-GAL4* might affect immune function and infection resistance. To test this prediction, we first systemically challenged flies carrying *yolk-GAL4* with the Gram-negative bacterial pathogen *Providencia rettgeri* (Juneja and Lazzaro 2009) and monitored population survival over five days post-infection. Strikingly, we found that females carrying *P{yolk-GAL4}2* were extremely sensitive to *P. rettgeri*. While only ∼50% of non-transgenic Canton-S flies died over the course of five days, the entire population of *P{yolk-GAL4}2* heterozygotes and homozygotes succumbed primarily within 24 hours and no later than 48 hours after administering *P. rettgeri* (Fig.4A). To determine whether *yolk-GAL4* females are particularly sensitive to *P. rettgeri* or whether the transgene increases susceptibility to bacterial infections in general, we also infected *yolk-GAL4*-expressing flies with the Gram-positive pathogen *Enterococcus faecalis* (Lazzaro 2002) and with the Gram-negative bacterium *Pectinobacter* (*Erwinia*) *carotovora* (*Ecc15*; Basset et al. 2000), which has relatively low virulence to wild-type flies (Troha and Im et al. 2018). Similarly to *P. rettgeri* infection, we found that >97% of *yolk-GAL4*-expressing females succumbed to *E. faecalis* infection within 48 hours (Fig.4B). Approximately 75% of females homozygous for *P{yolk-GAL4}2* succumbed to *Ecc15* infection, whereas Canton-S flies experienced ∼10% population mortality (Fig.4C). We also observed a notable transgene dose effect. Although mortality levels for *P{yolk-GAL4}2/+* and *P{yolk-GAL4}2 /P{yolk-GAL4}2* females infected with *E. faecalis* were comparable by five days post-infection, homozygotes succumbed more rapidly than heterozygotes (Fig.4B). The lesser virulence of *Ecc15* revealed a particularly striking effect of transgene copy number: ∼75% of *P{yolk-GAL4}2* homozygous females succumbed to *Ecc15* infection, in contrast to the ∼34% mortality of *P{yolk-GAL4}2* heterozygotes (Fig.4C). Importantly, 100% of heterozygous and homozygous *yolk-GAL4* females survived septic injury with sterile PBS (n=51-89 flies per genotype), indicating that the thoracic wounding procedure alone does not contribute to the infection sensitivity of *yolk-GAL4* flies.

**Figure 4.**
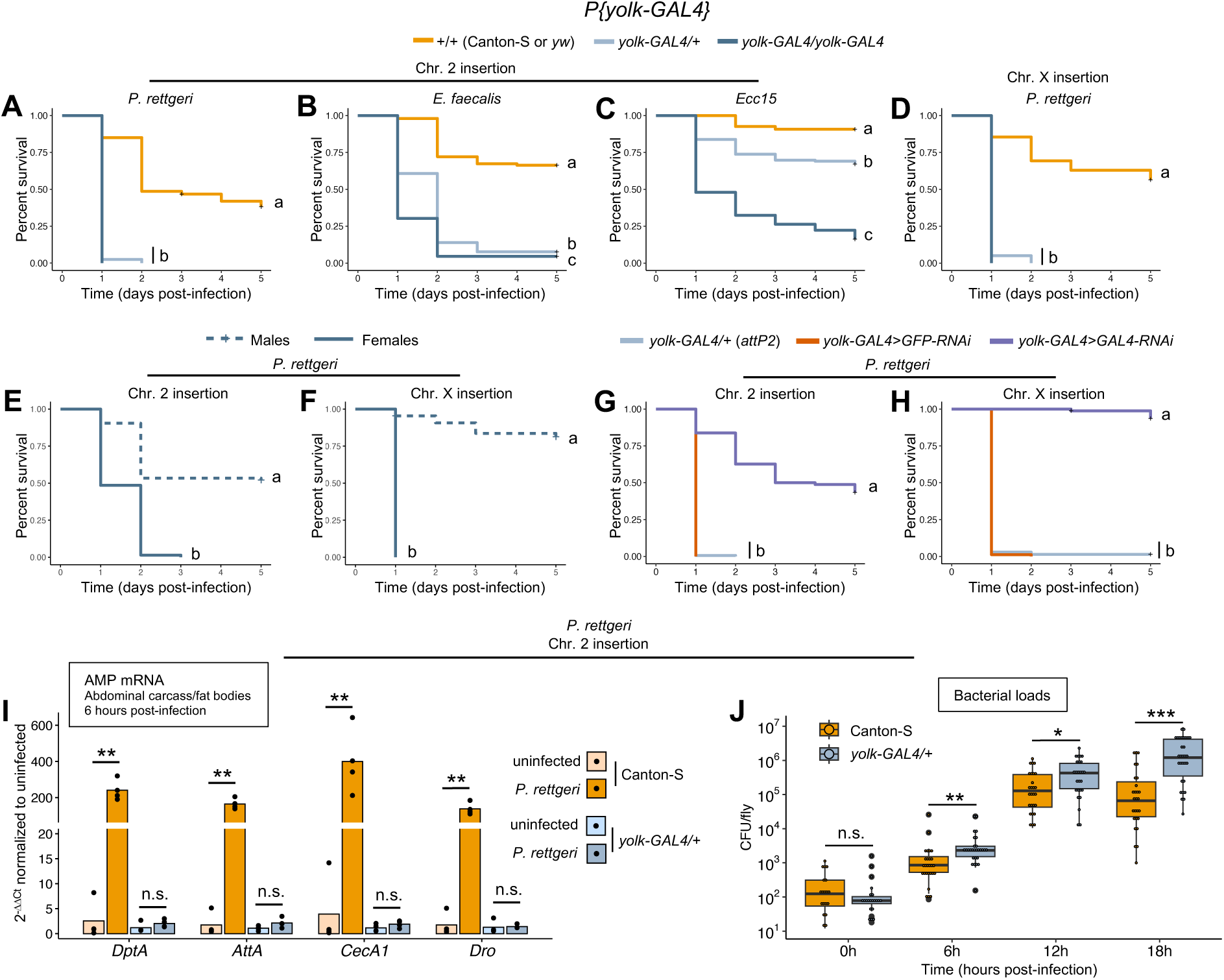
Expression of *yolk-GAL4* compromises resistance to systemic bacterial infections. (**A-C**) Females heterozygous or homozygous for the second chromosome *yolk-GAL4* insertion rapidly succumb to systemic infection with *Providencia rettgeri* **(A)** and *Enterococcus faecalis* **(B)**, and are significantly more susceptible to infection with *Pectobacterium carotovorum* (*Ecc15*; **C**). **(D)** Heterozygosity or homozygosity for the X chromosome *yolk-GAL4* insertion also causes extreme sensitivity to *P. rettgeri* infection. In panels **(A-C)** the non-transgenic wild-type control genotype (+/+; orange line) is Canton-S and in panel **(D)** the control genotype is *yw*. **(E,F)** Male flies which carry but do not express (see Fig.S1H,H’) either the second chromosome **(E)** or X chromosome **(F)** *yolk-GAL4* insertion survive *P. rettgeri* infection similarly to non-transgenic flies. **(G,H)** Knocking down expression of either the second chromosome **(G)** or X chromosome **(H)** *yolk-GAL4* insertion by RNAi fully restores females’ ability to survive *P. rettgeri* infection. **(I)** Females heterozygous for *yolk-GAL4* fail to upregulate expression of genes coding for antimicrobial peptides following *P. rettgeri* infection. RT-qPCR analysis of abdominal carcasses/fat bodies dissected from unchallenged females or females 6 hours post-infection. Data represent 2^-11Ct^ values normalized to the mean ΔC_t_ value for uninfected samples within each genotype. Each dot represents an individual sample of 10 pooled fat bodies/abdominal carcasses, bars represent the mean. **(J)** Females heterozygous for *yolk-GAL4* sustain higher bacterial loads than non-transgenic control flies over the initial stages of infection with *P. rettgeri*. Each dot represents the bacterial load of an individual fly. Statistics: Infection survival data **(A-H)** were analyzed by pairwise Log-Rank tests with Benjamini Hochberg correction for multiple comparisons. Genotypes with different letters are statistically different from one another (p<0.05). RT-qPCR data **(I)** were analyzed by unpaired two-sample t-tests. Bacterial load data **(J)** were analyzed by Mann-Whitney test (0h), or by unpaired two-sample tests (6h, 12, 18h). *p<0.05, **p<0.01, ***p<0.001, n.s.=not significant.

We conducted several additional experiments to confirm that infection susceptibility is caused by *GAL4* expression and not by confounding effects of the transgene insertion. First, we confirmed that X-chromosome-inserted *P{yolk-GAL4}1* heterozygotes and homozygotes also exhibited complete population mortality within 24-48 hours post-infection with *P. rettgeri* (Fig.4D), matching the mortality observed in *P{yolk-GAL4}2* females. Next, we infected *P{yolk-GAL4}1* and *P{yolk-GAL4}2* males, reasoning that these flies are genetically identical to females (excepting X/Y chromosome copy number) but they do not express the transgene due to the sex-specific activity of the *yolk* enhancer (Fig.S1H,H’). In contrast to their female siblings, males of both genotypes survived *P. rettgeri* infection at predicted wild-type levels (Fig.4E,F, comparable to female Canton-S and *yw* in Fig.4A,D). Lastly, we knocked down *GAL4* expression in *yolk-GAL4* heterozygotes by RNAi. *GAL4* knockdown fully suppressed the infection susceptibility caused by *P{yolk-GAL4}2* (Fig.4G) and *P{yolk-GAL4}1* (Fig.4H). In summary, these data show that *yolk-GAL4* expression renders flies less able to survive systemic infections.

We next asked whether the infection susceptibility of *yolk-GAL4* females is indicative of an impaired humoral immune response. To assay immune function, we used RT-qPCR to quantify AMP expression levels in fat bodies dissected from Canton-S and *P{yolk-GAL4}2/+* females 6 hours post-infection with *P. rettgeri*. Concordant with the findings of Kim et al. (2014), we found that infection-induced AMP upregulation was severely attenuated in fat bodies of *P{yolk-GAL4}2* heterozygotes compared to the strong transcriptional induction observed in Canton-S fat bodies (Fig.4I). Females carrying *P{yolk-GAL4}2* also sustained significantly higher bacterial loads than wild-type animals at 6, 12, and 18 hours post-infection (Fig.4J), suggesting the weakened immune response resulted in failure to contain pathogen growth.

Taken together, these results show the *yolk-GAL4* transgene impairs humoral immune activation in the fat body, resulting in significantly compromised infection resistance.

### Fat body *GAL4* expression levels correlate with *P. rettgeri* infection susceptibility

Our results thus far show that *GAL4* expressed under control of the *yolk* regulatory sequence disrupts energy storage, oogenesis, and infection resistance. We hypothesized that some or all of these traits might be sensitive to *GAL4* expression strength, and that higher levels of *GAL4* in the fat body would result in more severe functional disruption. As a strategy to address this, we presupposed that distinct fat body driver lines might exhibit varied *GAL4* expression levels and that we could use these to test the correlation between transgene expression intensity and fat body-related phenotypes. We therefore examined four additional *GAL4* lines commonly used for genetic manipulations in the fat body: *3.1Lsp2-GAL4*, *C564-GAL4*, *r^4^-GAL4*, and *Lpp-GAL4* (Table 1).

**Table 1.**
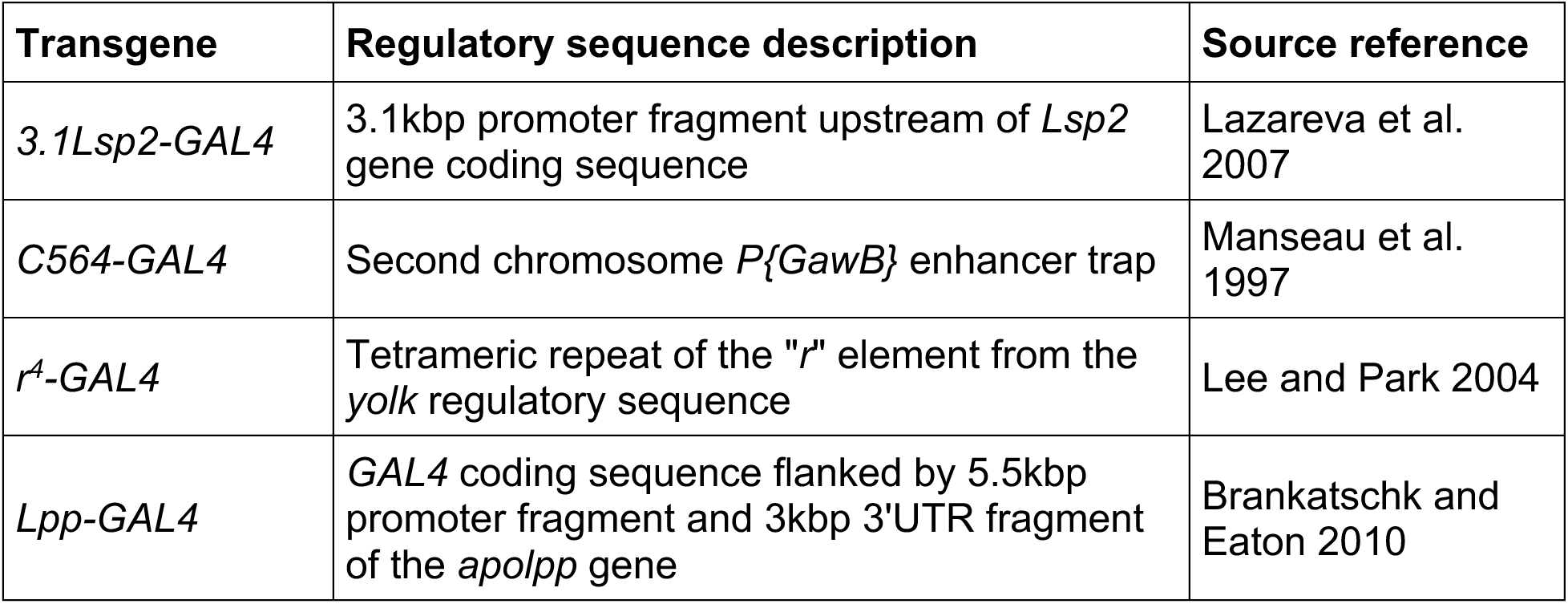
Fat body *GAL4* driver lines.

| Transgene | Regulatory sequence description | Source reference |
| --- | --- | --- |
| *3.1Lsp2-GAL4* | 3.1kbp promoter fragment upstream of *Lsp2* gene coding sequence | Lazareva et al. 2007 |
| *C564-GAL4* | Second chromosome *P{GawB}* enhancer trap | Manseau et al. 1997 |
| *r<sup>4</sup>-GAL4* | Tetrameric repeat of the "r" element from the *yolk* regulatory sequence | Lee and Park 2004 |
| *Lpp-GAL4* | *GAL4* coding sequence flanked by 5.5kbp promoter fragment and 3kbp 3'UTR fragment of the *apolpp* gene | Brankatschk and Eaton 2010 |

To confirm that the drivers we selected are all expressed in the adult fat body and to assess their tissue specificity, we first crossed each line to a *UAS-mCD8::GFP* reporter and visualized their expression patterns across major adult tissues in young adult females (Fig.S4). As expected, all four lines induced GFP expression in the abdominal fat body (Fig.S4Ba,Ca,Da,Ea). Notably, *3.1Lsp2-GAL4* produced extremely faint, almost undetectable live fluorescent signal at 25°C (Fig.S4Aa), but drove a mosaic pattern of fat body GFP expression when flies were housed at 29°C (Fig.S4Ba), which increases GAL4 activity (Duffy 2002). Consistent with prior reports (Lazareva et al. 2007; Brankatschk and Eaton 2010; Armstrong et al. 2014), *3.1Lsp2-GAL4* and *Lpp-GAL4* induced highly specific fat body expression (Fig.S4Ba,Ea), with no visible GFP fluorescence in the brain, flight muscle, gut, Malpighian tubules, or ovaries (Fig.S4Ab-Af,Bb-Bf,Eb-Ef). In addition to the fat body (Fig.S4Ca,Da), *C564-GAL4* and *r^4^-GAL4* both produced regionalized GFP expression in the gut (Fig.S4Cd,Ce,Dd,De), though not in other tissues (Fig.S4Cb,Cc,Cf,Db,Dc,Df). *C564-GAL4* activated GFP expression in the anterior midgut (Fig.S4Cd), and a middle midgut region which, based on its position and morphology, we suspect spans the stomach-like R3 compartment (Fig.S4Ce; Buchon et al. 2013). The *r^4^-GAL4* driver activated GFP expression in the proventriculus and crop duct (Fig.S4Dd), and throughout the hindgut (Fig.S4De).

**Figure S4.**
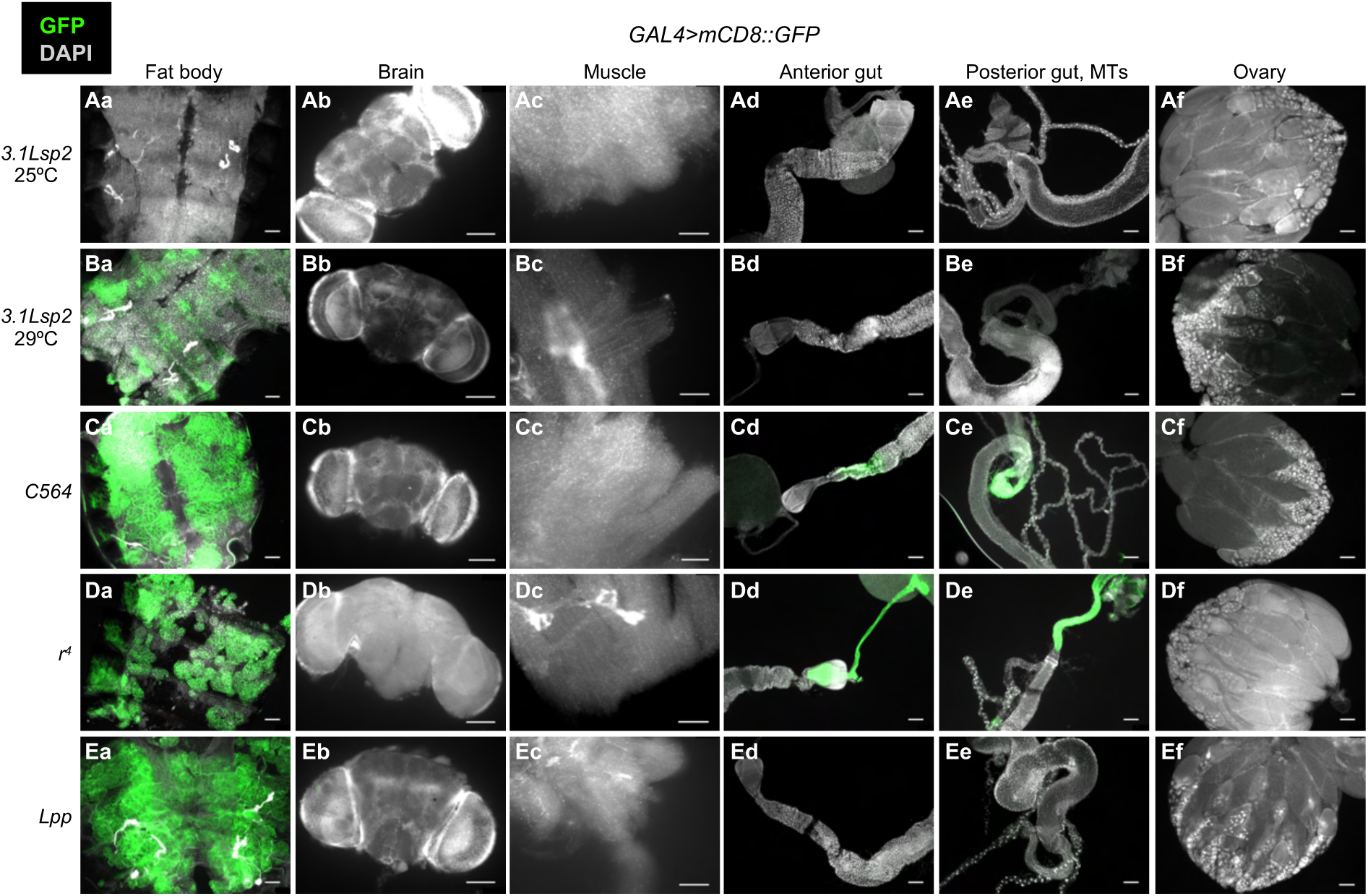
Expression patterns of fat body *GAL4* transgenes in adult female tissues. Flies carrying the indicated *GAL4* transgene were crossed to a *UAS-mCD8::GFP* reporter line. Images represent tissues dissected from F1 progeny to assess live GFP expression (green) induced by each driver. **(Aa-Bf)** *3.1Lsp2-GAL4* activates visible GFP expression specifically in the fat body only when flies are maintained at 29°C **(Aa** *vs.* **Ba)**. **(Ca-Cf)** *C564-GAL4* activates GFP expression in the fat body **(Ca)** and also in the anterior **(Cd)** and middle-posterior midgut **(Ce)**. **(Da-Df)** *r^4^-GAL4* activates GFP expression in the fat body **(Da)**, the crop duct and proventriculus **(Dd)**, and the hindgut **(De)**. **(Ea-Ef)** *Lpp-GAL4* activates strong GFP expression specifically in the fat body **(Ea)**. Nuclei are stained with DAPI (grey) to delineate overall tissue morphology. Scale bars=100µm.

After confirming each driver’s expression pattern, we next directly measured transgene expression strength by quantifying *GAL4* transcript levels in the fat body. To accomplish this, we crossed *3.1Lsp2-GAL4*, *C564-GAL4*, *r^4^-GAL4*, *Lpp-*GAL4, or *yolk-GAL4* to Canton-S flies, and used RT-qPCR to quantify *GAL4* expression in fat bodies dissected from the resulting female progeny heterozygous for each transgene. As we anticipated, this analysis revealed considerable variation in fat body *GAL4* transcript levels across the five lines. Consistent with our imaging results (Fig.S4Aa,Ba), we found that *3.1Lsp2-GAL4* flies expressed *GAL4* at the lowest levels relative to the other four lines (Fig.5A). Relative to *3.1Lsp2-GAL4* as a baseline, *GAL4* expression was, on average, 8-fold higher in fat bodies of *C564-GAL4* flies and 238-fold higher in *r^4^-GAL4* flies (Fig.5A). *Lpp-GAL4* and *yolk-GAL4* displayed the strongest fat body *GAL4* expression, with both exhibiting >2900-fold higher mRNA relative to *3.1Lsp2-GAL4* (Fig.5A).

**Figure 5.**
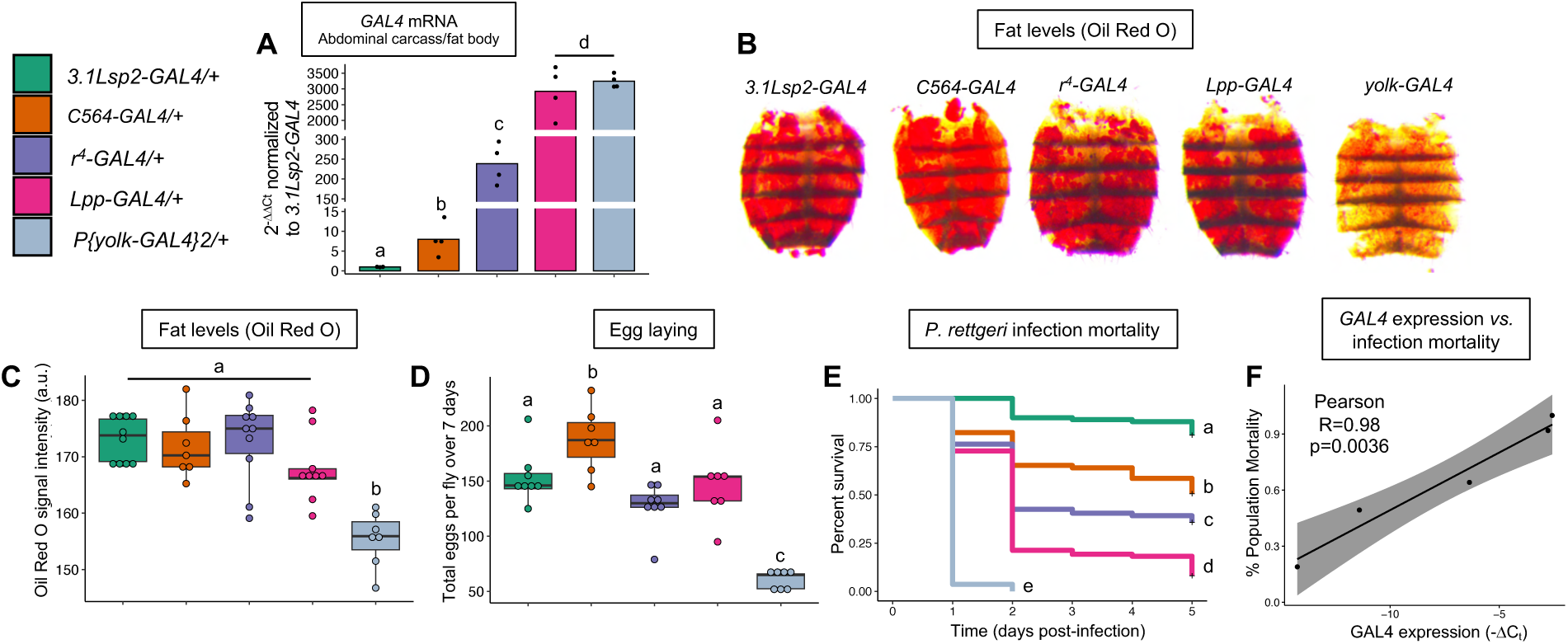
*GAL4* expression in the adult fat body correlates with *P. rettgeri* infection sensitivity. **(A)** RT-qPCR analysis documents a range of *GAL4* expression levels in abdominal carcasses dissected from females heterozygous for different fat body driver transgenes. Data represent 2^-11Ct^ values normalized to the mean ΔC_t_ value for *3.1Lsp2-GAL4/+* flies. Each dot represents an individual sample of 10 pooled fat bodies/abdominal carcasses, bars represent the mean. **(B)** Representative images of Oil Red O-stained abdominal carcasses dissected from females heterozygous for each of the indicated *GAL4* transgenes. **(C)** Quantification of Oil Red O staining intensities indicates that only *yolk-GAL4* results in decreased abdominal lipid levels while the other drivers do not differ in fat content. Each dot represents an individual fly/abdomen, a.u.=arbitrary units. **(D)** Additional fat body driver transgenes do not result in reduced egg laying rates comparable to *yolk-GAL4*. Each dot represents an individual replicate, 2-3 females per replicate. **(E,F)** Across the five *GAL4* transgenes assayed, mortality induced by *P. rettgeri* infection **(E)** directly correlates with *GAL4* expression levels in the fat body **(F)**. The correlation analysis in **(F)** plots the percentage of flies that succumbed by 5 days post-infection (y-axis; from data represented in E) against *GAL4* expression levels (x-axis; from data represented in A), quantified as the negative average ΔC_t_ value (Ct_GAL4_-Ct_Rpl32_; increasing value indicates higher *GAL4* expression). Statistics: Oil Red O signal intensity data **(C)** and egg laying data **(D)** were analyzed by one-way ANOVA with Tukey’s post hoc pairwise comparisons. Infection survival data **(E)** were analyzed by pairwise Log-Rank test with Benjamini Hochberg correction for multiple comparisons. Genotypes with different letters are statistically different from one another (p<0.05).

Having documented a wide range of *GAL4* transcript levels across our five selected drivers and noting that *yolk-GAL4* was one of the highest-expressing lines, we leveraged this variation to ask whether *GAL4* expression strength determines the severity of impacts on fat body-regulated traits. We measured fat levels, egg laying, and mortality from *P. rettgeri* infection in females carrying each of the five *GAL4* transgenes, again examining heterozygous progeny from crosses to Canton-S.

We found no significant correlation between *GAL4* expression levels and fat levels or egg output across driver lines. ORO lipid staining intensities were not significantly different among flies heterozygous for *3.1Lsp2-GAL4/+*, *C564-GAL4/+*, *r^4^-GAL4/+*, and *Lpp-GAL4/+*, although all were higher than *yolk-GAL4/+* females (Fig.5B,C). Similarly, egg output was comparable among females carrying *3.1Lsp2-GAL4*, *r^4^-GAL4*, and *Lpp-GAL4*, with *C564-GAL4/+* females laying more eggs than any other line, and *yolk-GAL4/+* females again producing relatively low numbers of eggs (Fig.5D). However, we observed a striking correlation between *GAL4* expression levels and susceptibility to *P. rettgeri* infection (Fig.5E,F). Flies heterozygous for *3.1Lsp2-GAL4*, which express the lowest *GAL4* levels (Fig.5A), were the least susceptible to infection, with only ∼15% of the total population succumbing. *C564-GAL4/+* and *r^4^-GAL4/+* flies displayed intermediate susceptibility, with ∼40% and ∼50% death, respectively (Fig.5E). *Lpp-GAL4*, which is expressed comparably to *yolk-GAL4* (Fig.5A), caused relatively high mortality (∼80%), though not as severe as the 100% lethality of *yolk-GAL4/+* females (Fig.4A,5E). As with *yolk-GAL4*, we found that knocking down *GAL4* expression in *Lpp-GAL4* heterozygotes (Fig.S5A) significantly increased their ability to survive infection (Fig.S5B), confirming that *GAL4* is responsible for the susceptibility of this line.

**Figure S5.**
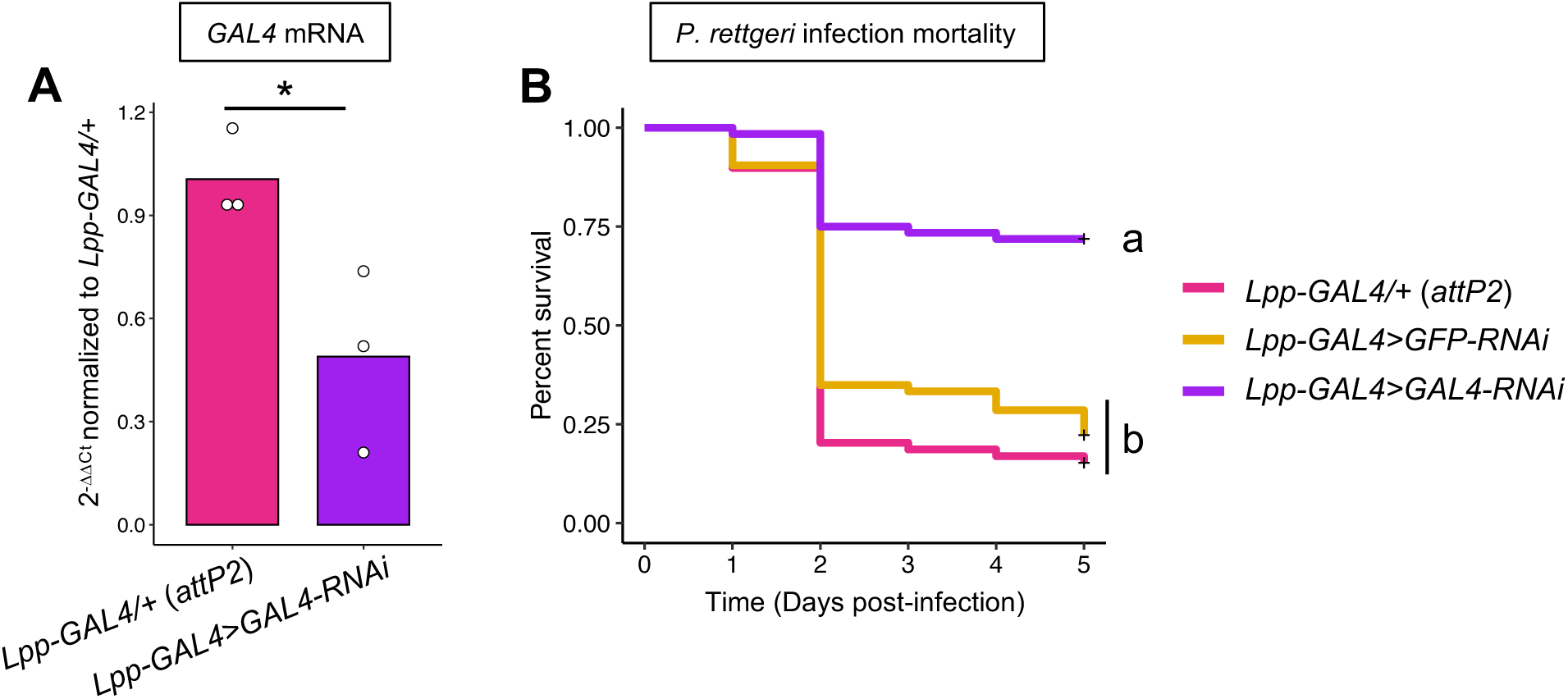
*GAL4* expression under control of the *Lpp* regulatory sequence increases infection sensitivity. **(A)** RT-qPCR analysis of whole flies indicates that activating the *UAS-GAL4-RNAi* construct with *Lpp-GAL4* reduces *GAL4* expression levels. Data represent 2^-11Ct^ values normalized to the mean ΔC_t_ value for control females with *Lpp-GAL4* heterozygous over the *attP2* genetic background. Each dot represents an individual sample of 5-10 pooled flies, bars represent the mean. *p<0.05, unpaired two-sample t-test. **(B)** RNAi knockdown of *GAL4* expression substantially increases infection survival of females heterozygous for *Lpp-GAL4*, indicating that GAL4 increases infection sensitivity of females carrying this driver transgene. Infection survival data were analyzed by pairwise Log-Rank tests with Benjamini Hochberg correction for multiple comparisons. Genotypes with different letters are statistically different from one another (p<0.05).

In summary, our data show that sensitivity to *P. rettgeri* infection quantitatively scales with *GAL4* expression in the adult female fat body, but stored fat levels and egg production do not. Nevertheless, expression of *GAL4* in the fat body controlled by the *yolk* enhancer impacts all three traits, so in our subsequent experiments we sought to investigate potential mechanisms by which this might occur.

### No evidence for GAL4-induced apoptosis in the adult female fat body

High levels of GAL4 have been reported to induce apoptosis in eye imaginal discs (Kramer and Staveley 2003) and in ventral lateral neurons of the adult brain (Rezával et al. 2007). We therefore hypothesized that overexpression of GAL4 might induce cell death in the fat body, which could contribute to the physiological dysfunction we documented in *yolk-GAL4* expressing flies. If the reduced infection resistance, egg production, and fat storage phenotypes caused by *yolk-GAL4* were due to fat body cell death, we reasoned that genetically inhibiting apoptosis should suppress these phenotypes. To test this, we crossed *yolk-GAL4* with flies carrying a *UAS* transgene encoding Baculovirus p35 protein, which potently inhibits apoptosis-activating caspases Dcp1 and Death related ICE-like caspase (Drice; Fig.S6A; Hay et al. 1994; Hawkins et al. 2000; Meier et al. 2000; Lannan et al. 2007). Compared to control animals expressing a non-functional *UAS-eGFP* transgene, we found that p35 expression did not increase fat levels (Fig.S6B,B’,C), egg output (Fig.S6D), or infection resistance (Fig.S6E) in *yolk-GAL4* heterozygotes. In fact, *yolk-GAL4>p35* females displayed a modest decrease in egg production relative to *yolk-GAL4>eGFP* controls (Fig.S6D). Successful completion of oogenesis requires apoptotic elimination of the follicle cells in late-stage oocytes (Nezis et al. 2002). Inhibition of this process by p35 might exacerbate the oogenesis defects in *yolk-GAL4* females. These results do not support the hypothesis that GAL4-induced apoptosis contributes to the deleterious phenotypes of *yolk-GAL4*-expressing flies.

**Figure S6.**
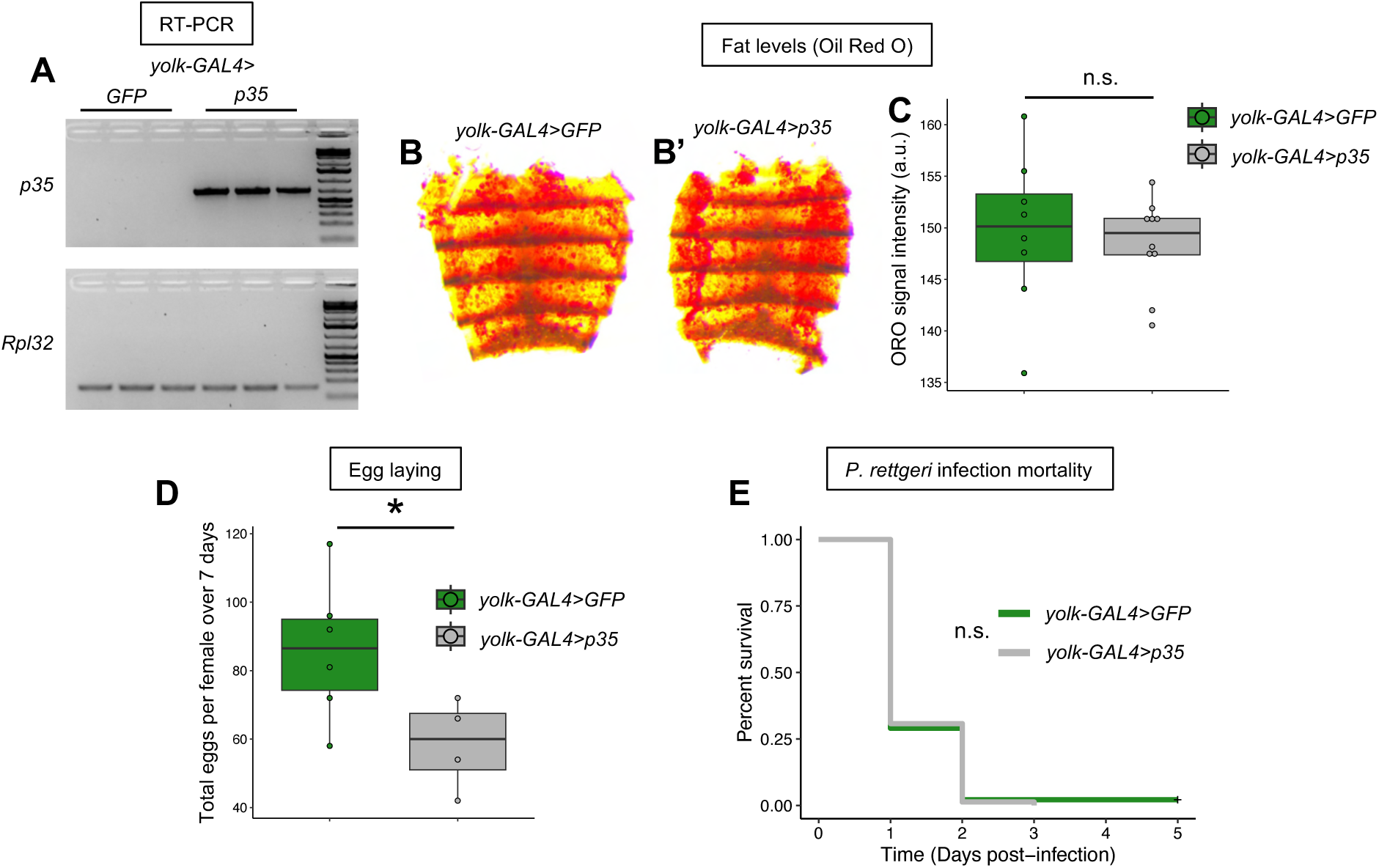
Inhibiting apoptosis does not suppress physiological defects caused by *yolk-GAL4* expression. **(A)** RT-PCR confirms *UAS-*activated expression of the apoptosis-inhibiting Baculovirus *p35* transgene induced by *yolk-GAL4*. No *p35* expression is detected in control animals with *yolk-GAL4* activating expression of a non-functional *UAS-GFP* transgene. The ubiquitously expressed *Rpl32* gene serves as a loading control. Each lane represents a separate sample of 5 pooled flies. **(B-C)** Expression of *p35* does not increase abdominal lipid content in females carrying *yolk-GAL4*. Representative images show abdominal ORO staining intensity appears similar between *yolk-GAL4>GFP* controls **(B)** and *yolk-GAL4>p35* flies **(B’’)**. Quantification of ORO signal indicates p35 has no effect on lipid levels **(C)**. Each dot represents an individual fly abdomen, a.u.=arbitrary units. **(D)** Females carrying *yolk-GAL4* and expressing *p35* lay fewer eggs than *yolk-GAL4>GFP* controls. Each dot represents an individual replicate, 2-3 females per replicate. **(E)** Expressing *p35* does not suppress the sensitivity to *P. rettgeri* infection caused by *yolk-GAL4*. Statistics: ORO data **(C)** and egg laying data **(D)** were analyzed by unpaired, two-sample t-test **(C,D)**. Infection survival data **(E)** were analyzed by Log-Rank test. *p<0.05, n.s.=not significant.

### Expressing nuclear-localized mCherry in the fat body reduces infection survival

We next asked whether high-level expression of transgenic proteins in general can affect fat body function, or whether the metabolic, reproductive, and immune phenotypes of *yolk-GAL4*-expressing flies are specifically caused by the GAL4 protein. We formulated two distinct hypotheses to explain how strong transgene expression might generally impact the fat body’s ability to execute key cellular and physiological functions. H1: Because the GAL4 transcription factor is actively imported into the nucleus (Uv et al. 2000), we hypothesized that import and/or accumulation of high levels of foreign protein in the nucleus might disrupt essential cellular processes like chromatin organization or gene expression, which would be consistent with the reduced expression of *Yolk Protein* (Fig.3G) and AMP genes (Fig.4I) we observed in *yolk-GAL4/+* fat bodies. H2: As a non-exclusive alternative, we hypothesized that expressing high levels of transgenic protein might exhaust cellular resources and overwhelm endogenous cellular machinery, thus impairing the fat body’s capacity to perform critical functions. We reasoned that if either of these proposed mechanisms were responsible for the defects we observed in *yolk-GAL4*-expressing flies, then high-level transgenic expression of other nuclear-localized proteins might elicit similar phenotypes. However, the two hypotheses can be distinguished. If general cellular strain caused by high-level expression of transgenic products (H2) impairs the fat body’s metabolic, reproductive, or immune functions, then expressing either a cytoplasmic or a nuclear-localized transgenic protein in the fat body would result in phenotypes similar to those of *yolk-GAL4*. Alternatively, if nuclear import and/or accumulation of the transgenic protein (H1) primarily affects these physiological traits, we would expect other nuclear-localized transgenic proteins to phenocopy *yolk-GAL4* but the same transgenic protein would have no effect if it remained localized to the cytoplasm.

To resolve these alternatives, we generated two new transgenic fly lines in which the *yolk* enhancer sequence directs expression of the fluorescent protein mCherry. The two lines are identical except one construct includes N– and C-terminal nuclear localization signal sequences (*yolk-mCherry.nls*), while the other lacks these sequences (*yolk-mCherry.cyt*). Both *yolk-mCherry.nls* and *yolk-mCherry.cyt* were integrated at the same *attP40* docking site on the second chromosome, enabling us to directly compare the effects of nuclear import *versus* protein synthesis on metabolic, reproductive, and immune physiology. We selected mCherry as an alternative transgenic protein to test our hypotheses for two reasons. First, mCherry allowed us to easily assess expression and subcellular localization of the transgenic proteins by fluorescence microscopy. Second, in the event that fly physiology was not strongly affected, we reasoned that these lines would constitute novel transcriptional reporters broadly useful for *Drosophila* researchers studying mechanisms of enhancer activation and *yolk* gene regulation.

To confirm their expression patterns, we first visually examined mCherry fluorescence in tissues dissected from *yolk-mCherry.nls* and *yolk-mCherry.cyt* flies (Fig.6A-D’, Fig.S7). As expected, both lines exhibited robust mCherry signal throughout the adult female fat body (Fig.6A-B’, Fig.S7A,A’,G,G’), and in the ovarian follicle cells (Fig.6C-D’, Fig.S7B,B’,H,H’). Additionally, mCherry was strikingly nuclear-localized in fat bodies and ovaries of *yolk-mCherry.nls* females (Fig.6A,C, Fig.S7A,B), in contrast to the more diffuse fluorescence observed in *yolk-mCherry.cyt* tissues (Fig.6B,D, Fig.S7G,H). Notably, unlike *yolk-GAL4* (Fig.S1E,J), we did not observe mCherry signal in the crop duct for either line (Fig.S7E,E’,K,K’). We also did not detect fluorescence in the brain, gut, Malpighian tubules, or muscle (Fig.S7C-F’,I-L’), nor in the fat bodies of late third instar female larvae (Fig.S7M,M’,O,O’) or adult males (Fig.S7N,N’,P,P’). Thus, our *mCherry* transgenes largely recapitulate the spatiotemporal expression patterns of *yolk-GAL4* and display the expected subcellular localization of fluorescent protein.

**Figure 6.**
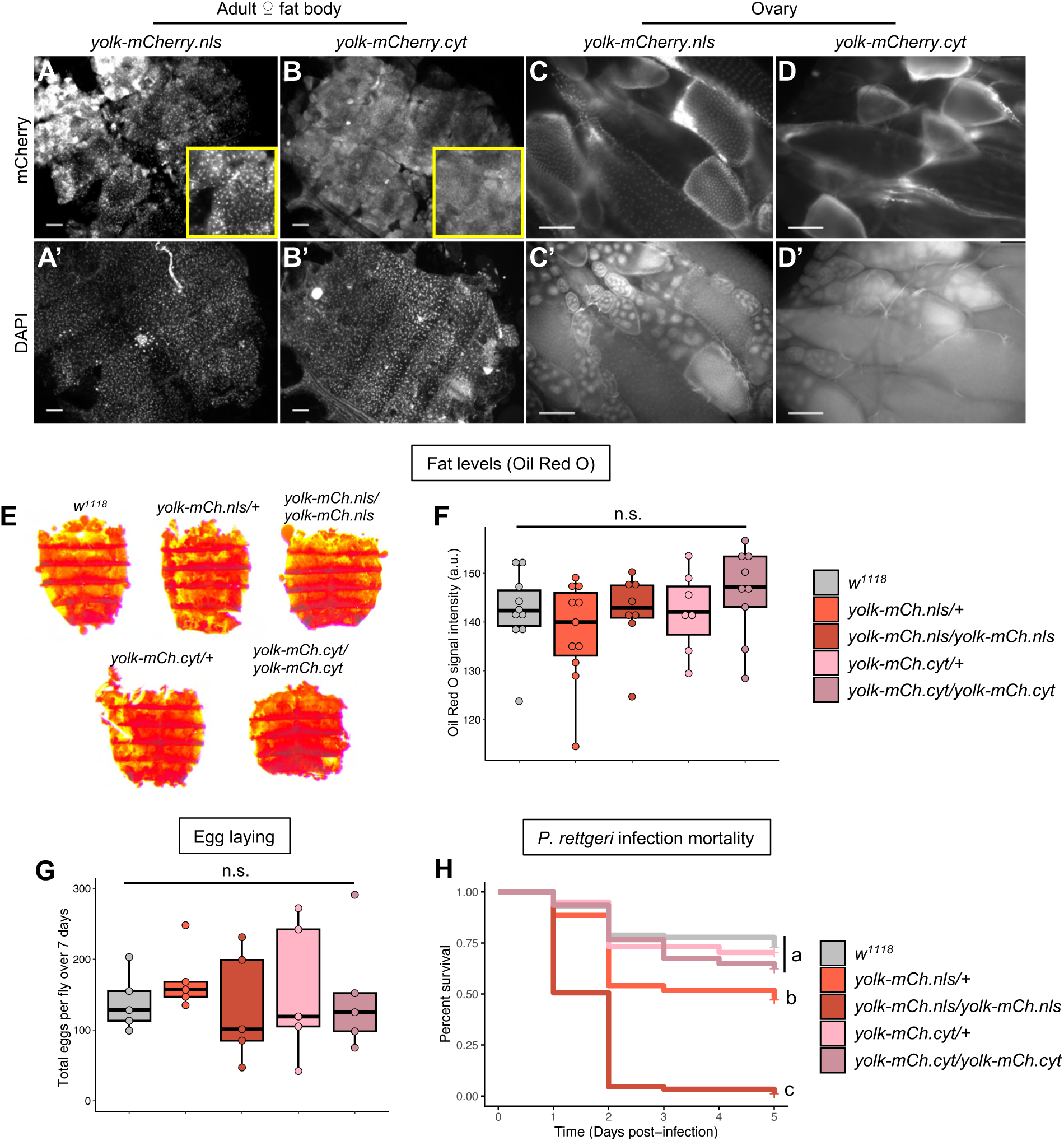
Nuclear-localized mCherry expressed in the fat body with the *yolk* enhancer reduces infection survival but not lipid levels or egg laying. (A-D’) The *yolk-mCherry* transgenes are expressed in the adult female fat body **(A-B’)** and in ovarian follicle cells **(C-D’)**. The *yolk-mCherry.nls* construct, with nuclear-localization sequences flanking mCherry, shows strongly nuclear-localized mCherry signal in the fat body and ovary **(A,C)**, while the *yolk-mCherry.cyt* transgene produces mCherry that remains localized in the cytoplasm **(B,D)**. Nuclei are stained with DAPI **(A’,B’,C’,D’)** to reveal overall tissue morphology. Scale bars=100µm. **(E,F)** Expressing cytoplasmic or nuclear-localized mCherry under control of the yolk enhancer does not affect abdominal lipid levels. **(E)** Representative images of Oil Red O-stained abdominal carcasses dissected from *w^1118^*control females or from females heterozygous or homozygous for either *yolk-mCherry.nls* or *yolk-mCherry.cyt*. **(F)** Quantification of ORO staining intensity. Each dot represents an individual fly/abdomen, a.u.=arbitrary units. **(G)** Females heterozygous or homozygous for either *yolk-mCherry* construct lay comparable numbers of eggs to non-transgenic control females. Each dot represents an individual replicate, 2-3 females per replicate. **(H)** Females expressing nuclear-localized mCherry in the fat body exhibit transgene dose-dependent increased mortality caused by *P. rettgeri*, with one copy of *yolk-mCherry.nls* moderately increasing infection sensitivity relative to *w^1118^* controls and *yolk-mCherry.nls* homozygotes exhibiting high susceptibility to infection. Cytoplasmically retained mCherry produced in the same expression pattern does not affect females’ susceptibility to *P. rettgeri* infection, with *yolk-mCherry.cyt* heterozygotes and homozygotes surviving comparably to *w^1118^* females. Statistics: ORO data **(F)** and egg laying data **(G)** were analyzed by one-way ANOVA. Infection survival data **(H)** were analyzed by pairwise Log-Rank test with Benjamini Hochberg correction for multiple comparisons. Genotypes with different letters are statistically different from one another (p<0.05), n.s.=not significant.

**Figure S7.**
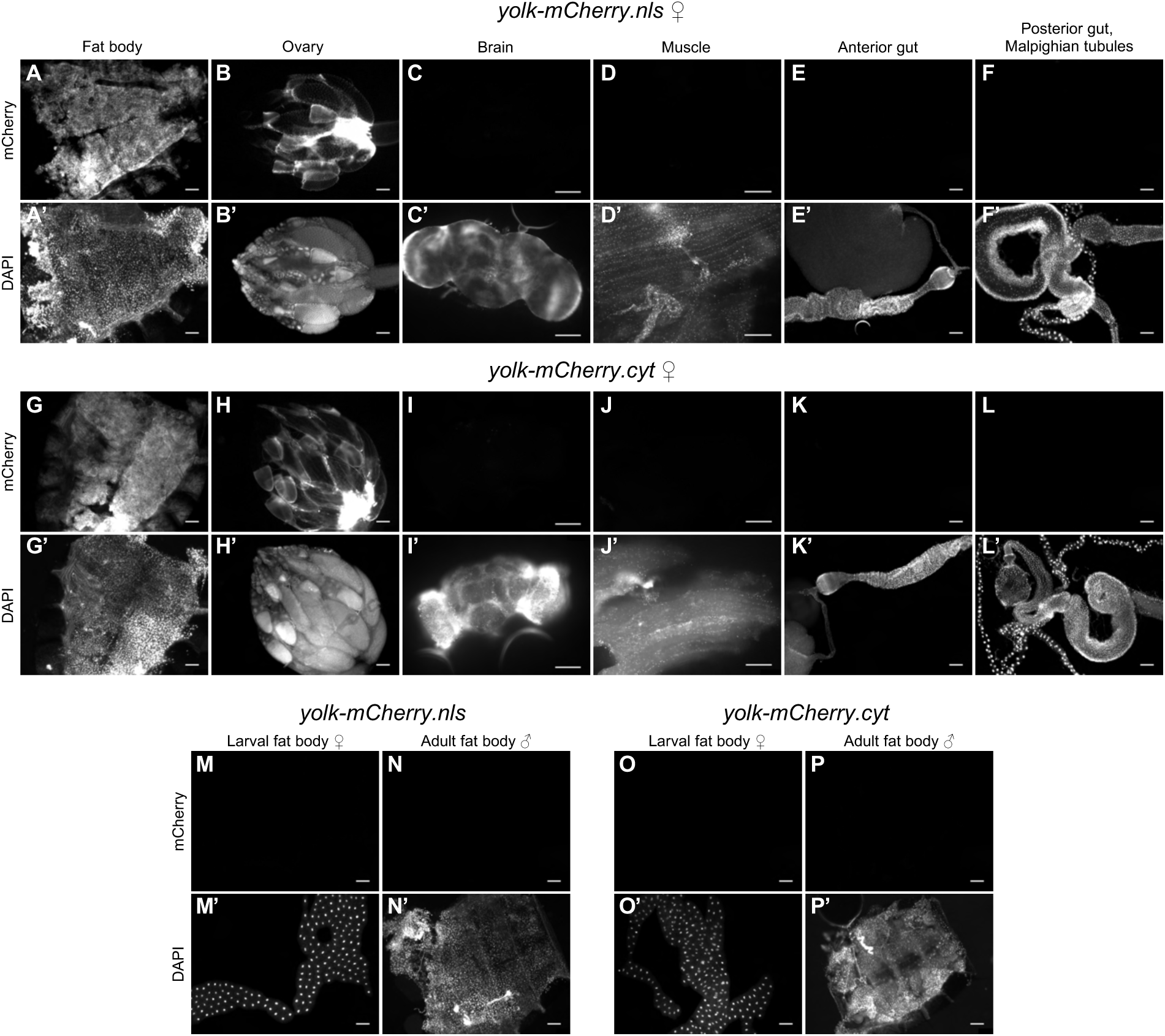
The *yolk-mCherry.nls* and *yolk-mCherry.cyt* transgenes recapitulate the *yolk-GAL4* expression pattern. Live mCherry fluorescence signal in major tissues dissected from adult females homozygous for the *yolk-mCherry.nls* **(A-F)** or yolk-*mCherry.cyt* **(G-L)** transgenes. For both constructs, strong mCherry signal is observable in the abdominal fat body **(A,A’,G,G’)** and in the follicle cells of maturing oocytes **(B,B’,H,H’)**. mCherry is not detectable in the brain **(C,C’,I,I’)**, muscle **(D,D’,J,J’)**, gut, or Malpighian tubules **(E-F’,K-L’)**. No mCherry fluorescence is visible in the fat bodies of larval females **(M,M’,O,O’)** or adult males **(N,N’,P,P’)**, indicating adult female-specific expression. Overall expression patterns of both constructs are therefore comparable to that of *yolk-GAL4* (see Fig.S1). Nuclei are stained with DAPI to delineate general tissue morphologies. Scale bars=100µm.

**Figure S8.**
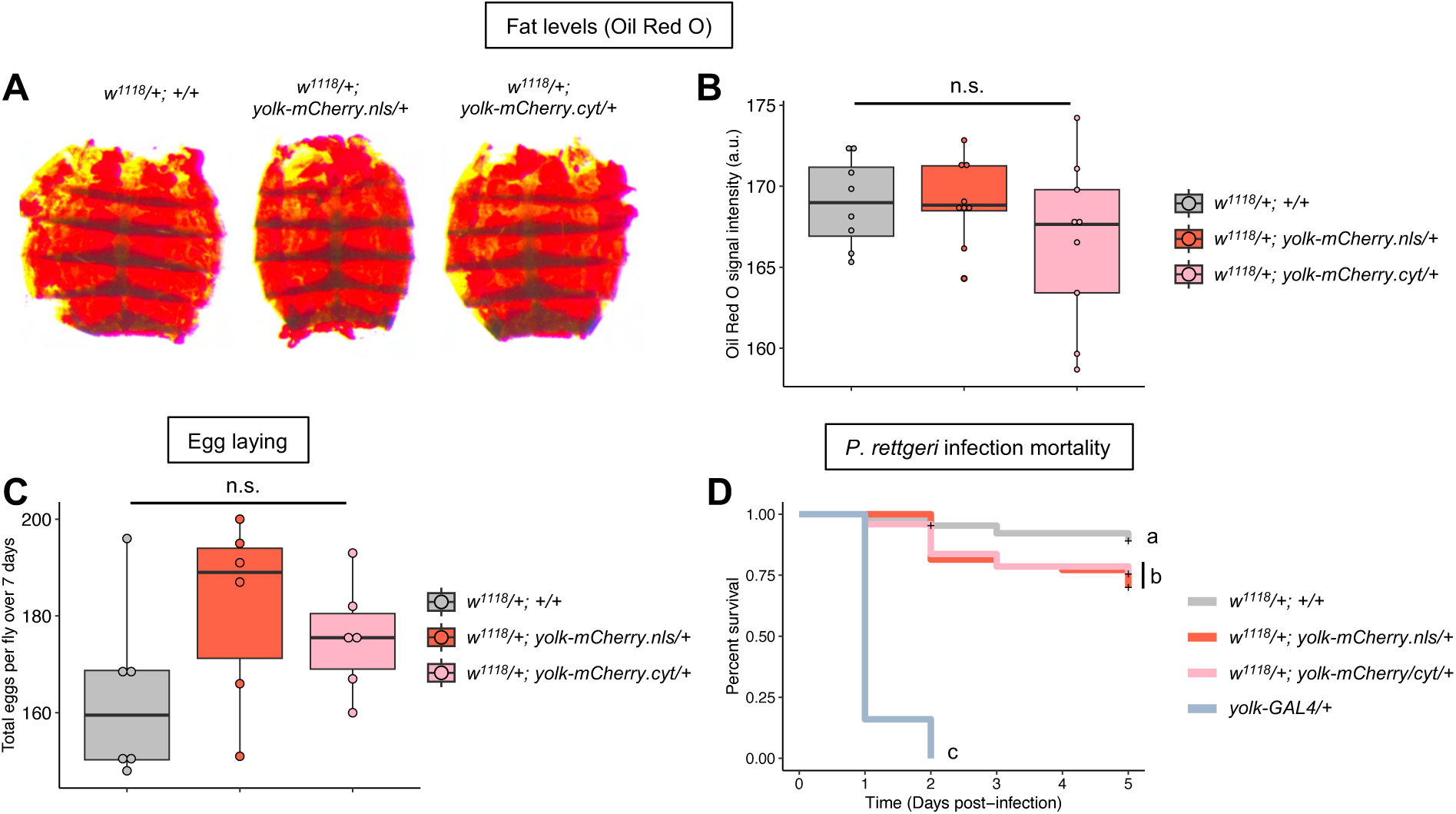
Heterozygosity for *yolk-mCherry.nls* or *yolk-mCherry.cyt* does not affect fat levels or egg laying, and minimally increases infection mortality. All genotypes represent F1 progeny of crosses between *w^1118^*, *yolk-mCherry.nls*, or *yolk-mCherry.cyt* males and Canton-S females. **(A,B)** Oil Red O staining of abdominal carcasses dissected from females heterozygous for either *yolk-mCherry.nls* or *yolk-mCherry.cyt* indicates comparable gross lipid levels between transgenic flies and non-transgenic *+/+* controls. In panel **(B)** each dot represents an individual fly/abdomen, a.u.=arbitrary units. **(C)** Females carrying one copy of either *yolk-mCherry.nls* or *yolk-mCherry.cyt* lay numbers of eggs comparable to those of non-transgenic controls. Each dot represents an individual replicate, 2-3 females per replicate. **(D)** Females heterozygous for either *yolk-mCherry.nls* or *yolk-mCherry.cyt* are moderately more susceptible to *P. rettgeri* infection than wild-type flies, but not display the extreme sensitivity of *yolk-GAL4* heterozygotes. Statistics: ORO signal intensity data **(B)** and egg laying data **(C)** were analyzed by one-way ANOVA. Infection survival data **(D)** were analyzed by pairwise Log-Rank test with Benjamini Hochberg correction for multiple comparisons. Genotypes with different letters are statistically different from one another (p<0.05), n.s.=not significant.

Having validated their expression, we used these new fly lines to directly ask whether overexpression and/or nuclear import of an alternative transgenic protein distinct from GAL4 might also impact metabolism, reproduction, or immunity. To accomplish this, we measured abdominal fat levels, egg laying rates, and survival of *P. rettgeri* infection in females heterozygous or homozygous for *yolk-mCherry.cyt* or *yolk-mCherry.nls*. Because *yolk-GAL4* produced strong phenotypes when heterozygous over a variety of genetic backgrounds, we examined *yolk-mCherry* heterozygotes both in *trans* to their *w^1118^* background and to the Canton-S genotype used for our prior analyses.

We found that fat body lipid levels and reproductive output were not affected by either mCherry transgene. Abdominal lipid staining with ORO appeared robust in females heterozygous or homozygous for *yolk-mCherry.nls* or *yolk-mCherry.cyt* and quantified staining intensities were statistically equivalent to those of non-transgenic control flies (Fig.6E,F, Fig.S8A,B). Similarly, females carrying *yolk-mCherry.nls* or *yolk-mCherry.cyt* did not display any reduction in egg laying compared to control females (Fig.6G, Fig.S8C). However, when challenged with *P. rettgeri*, we found that *yolk-mCherry.nls* reduced female’s capacity to survive infection in a transgene dose-dependent manner (Fig.6H, Fig.S8D). Females heterozygous for *yolk-mCherry.nls* over both *w^1118^* and Canton-S backgrounds displayed moderately but significantly increased infection-induced mortality (Fig.6H, Fig.S8D). By comparison, *yolk-mCherry.nls* homozygous females were highly susceptible to *P. rettgeri*, with >95% of the population succumbing by 24-48 hours post-infection (Fig.6H). In contrast, the *yolk-mCherry.cyt* transgene, producing the same mCherry protein localized to the cytoplasm, had no effect on infection mortality when heterozygous or homozygous compared to the *w^1118^* background control line (Fig.6H). When heterozygous over the Canton-S background, *yolk-mCherry.cyt* led to a small but significant reduction in infection survival, equivalent to that caused by *yolk-mCherry.nls* in *trans* to Canton-S (Fig.S8D). This suggests that *mCherry* expression alone might exert modest deleterious effects on infection susceptibility in certain genetic backgrounds, but nuclear localization of copious transgenic protein results in substantial sensitivity to infection.

In summary, these results show that expression of mCherry under control of the *yolk* enhancer does not by itself strongly impact fat-body-regulated physiological traits, but that nuclear import of mCherry dramatically increases susceptibility to infection without affecting fat stores or egg output. Our results therefore suggest that metabolism and oogenesis might be disrupted by *GAL4* expression specifically, while the ability to withstand infection might be more generally sensitive to nuclear-localized transgenic proteins expressed in the fat body.

### Conversion of *yolk-GAL4* into *yolk-LexA* alleviates physiological defects caused by *GAL4* expression

Given our observation that nuclear-localized GAL4 and mCherry reduced infection survival in a transgene dose-dependent manner (Fig.4B,C, Fig.5E, Fig.6H), we predicted that another nuclear-localized transgenic protein might similarly reduce infection survival without impacting fat levels or egg laying. To test this prediction, we applied Homology-Assisted CRISPR Knock-in (HACK; Lin and Potter 2016; Chang et al. 2022; Rankin et al. 2024) to *yolk-GAL4* flies in order to replace the *GAL4* coding sequence with a *LexA* transgene.

The HACK approach utilizes *in vivo* CRISPR gene editing to exchange *GAL4* for *LexA* in any extant driver line (Fig.7A, Fig.S9A).The *LexA* system is an alternative bipartite expression system commonly used in *Drosophila* in which the bacterial LexA transcription factor activates expression of responder elements downstream of its cognate *LexAop* target sequence (Lai and Lee 2006). For HACK-mediated conversion, genetic crosses place the target *GAL4* construct in *trans* to a donor cassette containing the “*LexA*” coding sequence, which comprises the *LexA* DNA binding domain fused to the *GAL4* activation domain (LexA::GAD, conventionally abbreviated to “LexA”; Lai and Lee 2006). The *LexA* donor sequence is flanked by homology arms to *GAL4* and preceded at the 5’ end by a *T2A* ribosomal skipping sequence. Guide RNAs targeting the *GAL4* coding sequence and a germline Cas9 source lead to directed cleavage of the *GAL4* transgene, and homology-directed repair replaces *GAL4* with the *LexA* cassette. The converted transgene thus comprises the same regulatory sequence of the original line (here, the *yolk* enhancer) directing expression of a truncated, N-terminal GAL4 fragment and the LexA transcription factor as a separate protein (Fig.7A; Lin and Potter 2016; Chang et al. 2022; Rankin et al. 2024). We reasoned that generating a *yolk-LexA* line by this approach would provide another tool to ask whether infection resistance can be broadly perturbed by nuclear-localized transgenic proteins. In addition, *yolk-LexA* would constitute a novel reagent useful for adult female fat body-specific genetic manipulations.

**Figure 7.**
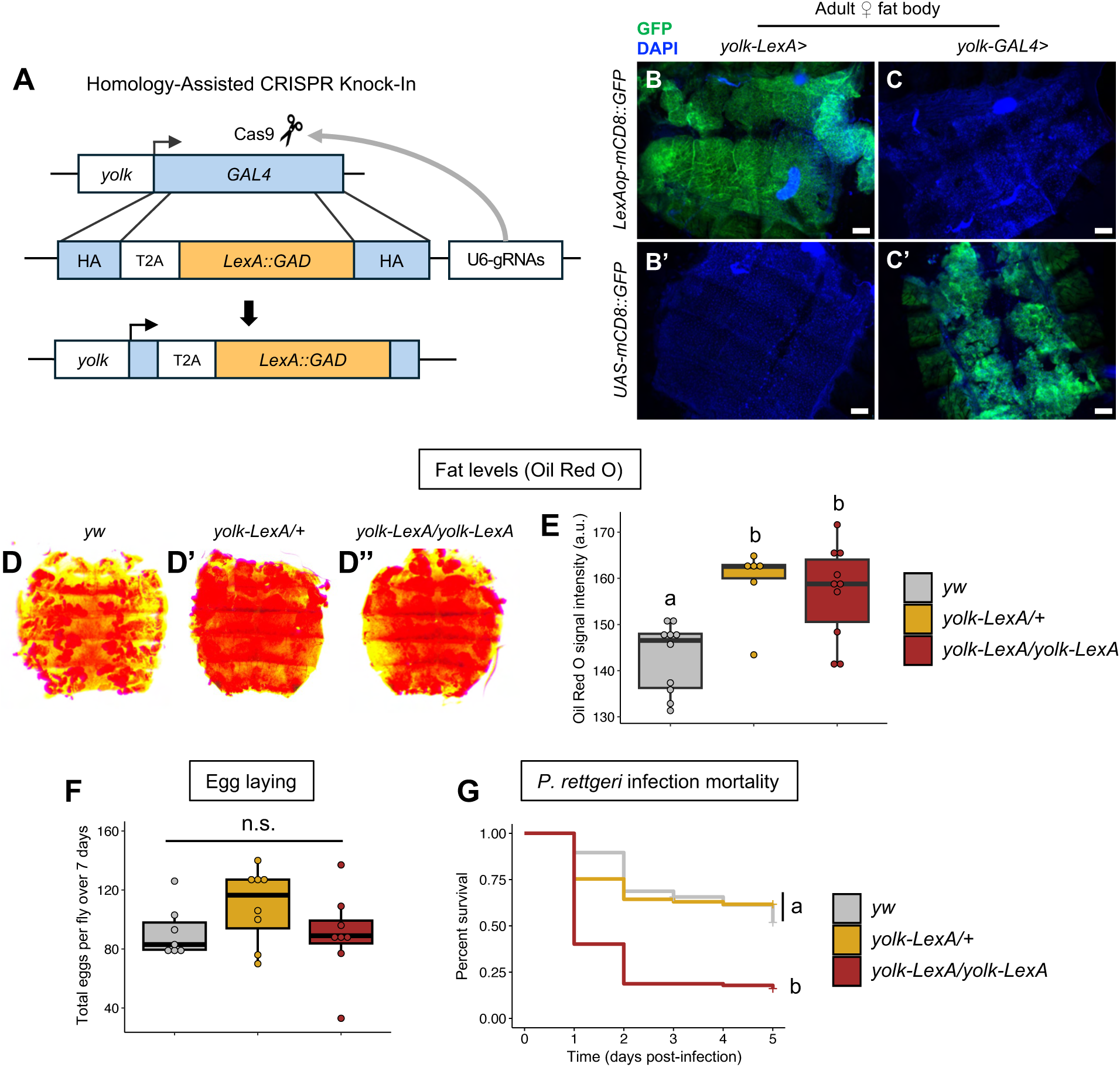
Converting *yolk-GAL4* to *yolk-LexA* alleviates metabolic and reproductive phenotypes, and lessens infection sensitivity caused by *GAL4* expression. (**A**) Schematic illustrating the HACK approach (Lin and Potter, 2016; Rankin et al., 2024) for *in vivo* CRISPR-mediated conversion of the second chromosome *yolk-GAL4* line into *yolk-LexA*. **(B-C’)** The converted *yolk-LexA* line activates expression of a *LexAop-mCD8::GFP* reporter in the adult female fat body **(B)** but does not activate expression of a *UAS-mCD8::GFP* reporter **(B’)**, indicating successful transgene conversion. The original *yolk-GAL4* transgene does not activate the *LexAop-mCD8::GFP* transgene **(C)** but does induce *UAS-mCD8::GFP* expression **(C’)**. Nuclei are labeled with DAPI to reveal general tissue morphology. Scale bars=100µm. **(D-E)** Unlike *yolk-GAL4* (Fig.2,S2), females carrying *yolk-LexA* do not display reduced abdominal lipid levels revealed by ORO staining. ORO signal is stronger in abdomens from flies with *yolk-LexA* than in *yw* background controls. In panel **(E)**, each dot represents an individual fly/abdomen, a.u.=arbitrary units. **(F)** Females heterozygous or homozygous for *yolk-LexA* lay numbers of eggs comparable to those of non-transgenic *yw* females. Each dot represents an individual replicate, 2-3 females per replicate. **(G)** Heterozygosity for *yolk-LexA* does not affect survival of *P. rettgeri* infection, while *yolk-LexA* homozygotes are more susceptible to infection. ORO data **(D)** were analyzed by Kruskal-Wallis test with Dunn’s post hoc pairwise comparisons. Egg laying data **(E)** were analyzed by one-way ANOVA. Infection survival data **(F)** were analyzed by pairwise Log-Rank tests with Benjamini Hochberg correction for multiple comparisons. Genotypes with different letters are statistically different from one another (p<0.05), n.s.=not significant.

Using a recently developed, efficient HACK donor (Rankin et al. 2024), we converted *yolk-GAL4* on the second chromosome into a *yolk-LexA* transgene. As expected, *yolk-LexA* drove robust GFP expression in the same pattern as *yolk-GAL4* when crossed to a *LexAop-mCD8::GFP* responder line (Fig.7B) but failed to activate *UAS-mCD8::GFP* (Fig.7B’), indicative of successful transgene conversion. This new line allowed us to ask whether an alternative, nuclear-localized transcription factor expressed from the identical genomic locus of the *P{yolk-GAL4}2* insertion would also affect fat levels, egg laying, or infection survival. We examined these traits in females heterozygous or homozygous for *yolk-LexA* in the *yw* genetic background in which they were constructed, and also in F1 heterozygotes from crosses with Canton-S.

We found that females heterozygous or homozygous for *yolk-LexA* did not exhibit reduced fat stores relative to non-transgenic controls (Fig.7D-E). In fact, abdominal ORO staining levels were higher in females carrying one or two copies of *yolk-LexA* (Fig.7D’,D’’) compared to their *yw* background line (Fig.7D,E). We also found that *yolk-LexA* had no effect on egg output, with heterozygotes and homozygotes laying total egg numbers comparable to those of *yw* females over seven days post-eclosion (Fig.7F). Similarly, females carrying *yolk-LexA* heterozygous over the Canton-S background displayed no reduction in abdominal fat levels (Fig.S9B,C) or egg production (Fig.S9D).

**Figure S9.**
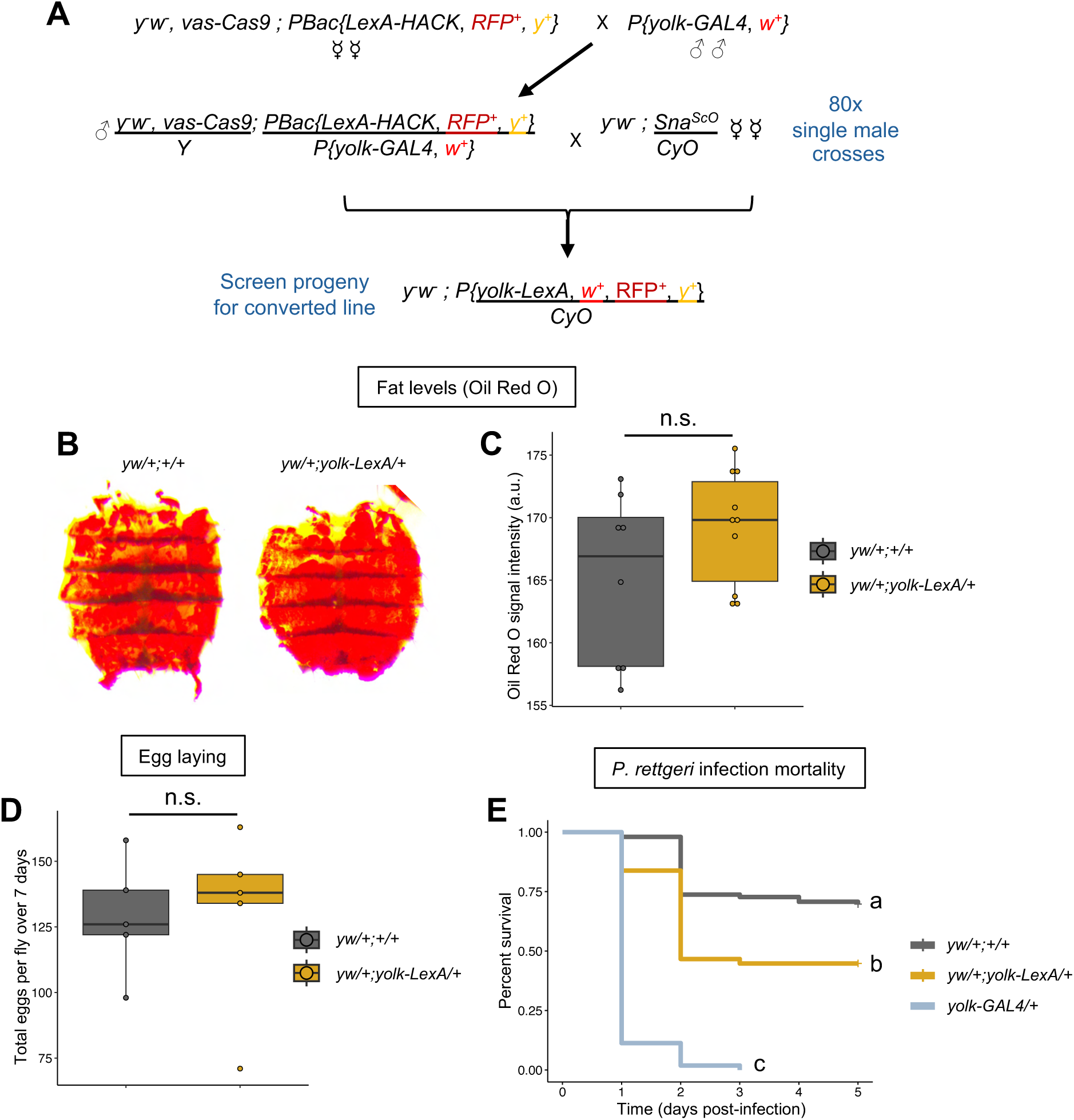
Heterozygosity for *yolk-LexA* does not affect fat levels or egg laying and moderately increases infection mortality. **(A)** Schematic adapted from Rankin et al. (2024) illustrating crossing scheme used to convert the second chromosome *P{yolk-GAL4}2* transgene to *yolk-LexA* using the v2 Homology-Assisted CRISPR Knock-in approach. **(B-E)** All genotypes represent F1 progeny of crosses between *yw* or *yolk-LexA* males and Canton-S females. **(B,C)** Females carrying one copy of the converted yolk-LexA transgene contain abdominal lipid levels comparable to those of non-transgenic control females. In **(C)** each dot represents an individual fly abdomen, a.u.=arbitrary units. **(D)** Egg laying rates are comparable between *yolk-LexA* heterozygotes and non-transgenic controls. Each dot represents an individual replicate, 2 flies per replicate. **(E)** Female *yolk-LexA* heterozygotes are more susceptible to *P. rettgeri* infection than non-transgenic control females but display substantially increased survival compared to females carrying one copy of the original *yolk-GAL4* transgene. Statistics: ORO data **(C)** and egg laying data **(D)** were analyzed by unpaired two-sample t-test. Infection survival data **(E)** were analyzed by pairwise Log-Rank tests with Benjamini Hochberg correction for multiple comparisons. Genotypes with different letters are statistically different from one another (p<0.05), n.s.=not significant.

In contrast to fat levels and egg laying, we observed a dose-dependent effect of *yolk-LexA* on females’ sensitivity to *P. rettgeri* infection, similar to that observed with *yolk-mCherry.nls*. One copy of *yolk-LexA* heterozygous over the *yw* genetic background had no effect on infection mortality (Fig.7G). When heterozygous over the Canton-S background, *yolk-LexA* led to significantly reduced infection survival compared to non-transgenic controls (Fig.S9E). However, ∼50% of these *yolk-LexA* heterozygotes were able to survive *P. rettgeri* infection, in striking contrast to the complete population mortality of *yolk-GAL4* heterozygotes (Fig.S9E). Notably, *yolk-LexA* homozygous females experienced substantially greater infection-induced mortality compared to heterozygous and control females (Fig.7G). Again, though, the infection sensitivity caused by two copies of *yolk-LexA* was less than that caused by a single copy of *yolk-GAL4*, as ∼15-20% of *yolk-LexA* homozygotes consistently survived infection (Fig.7G).

These data add further evidence that nuclear-localized transgenic proteins expressed at high levels in the adult fat body generally compromise the ability of *Drosophila* to survive pathogenic infection. Additionally, these results reinforce the conclusion that fat storage and egg production are more specifically affected by *GAL4* expression, as exchanging *GAL4* for *LexA* fully alleviates the metabolic and reproductive phenotypes caused by *yolk-GAL4*.

## DISCUSSION

The *GAL4-UAS* system is a mainstay for answering fundamental biological questions with *Drosophila*. However, the effects of transgenic *GAL4* expression and the GAL4 protein product itself on molecular, cellular, and physiological processes have only rarely been considered or empirically tested. Here we show that strong *GAL4* expression in the adult female fat body driven by the *yolk* regulatory sequence can severely compromise energy storage (Fig.2, Fig.S2), reproduction (Fig.3, Fig.S3), and immune performance (Fig.4). We find that enhancers which drive the transgene more weakly affect these traits to a lesser extent or not at all (Fig.5), indicating that the fat body is particularly sensitive to high levels of *GAL4* expression. Using several newly-generated transgenic lines, we present evidence that suggests the infection sensitivity of flies expressing *GAL4* and other transgenes is specifically caused by the localization of these proteins to fat body nuclei (Fig.6H, Fig.7G). Our work raises important considerations about the effects of transgenic tools on tissue homeostasis in *Drosophila* and other genetically amenable laboratory organisms, and about the limits on cellular function imposed by high levels of non-native proteins.

Several non-exclusive mechanisms might explain how high levels of transgenic GAL4 in fat body cells result in disrupted tissue integrity, and reduced lipid levels, egg production, and infection resistance. The fat body simultaneously executes biologically varied and energetically demanding processes, including synthesis and breakdown of macromolecular stores and secretion of large quantities of proteins, endocrine molecules, and metabolites (Canavoso et al. 2001; Arrese and Soulages 2010; Zheng et al. 2016). Allocation of a finite set of molecular resources among these processes can, in some cases, overburden the functional capacity of the fat. For example, mating-induced protein synthesis has been shown to constrain the fat body’s ability to mount an immune response, leading to activation of stress responses characteristic of overburdened translational capacity (Gupta et al. 2022). The demands of synthesizing large quantitates of GAL4 or other transgenic proteins at high levels might divert adipocyte resources from producing molecules required for metabolism, reproduction, and immunity. Additionally, high-level production of superfluous protein could overwhelm the functional capacities of core cellular machinery and organelles, leading to activation of stress responses that arrest overall tissue function. Notably, we only observed poor tissue integrity, reduced fat stores, and low egg output in females expressing *yolk-GAL4*, while flies expressing *GAL4* from other, comparatively weaker fat body promoters did not exhibit these defects (Fig.5B,C,D). It remains possible that GAL4 might impact fat body metabolic function at more nuanced levels not resolved by our low-resolution lipid staining analysis. Furthermore, our data suggest that oogenesis defects unique to *yolk-GAL4* flies are at least partially attributable to *GAL4* expression in follicle cells (Fig.S3G-I). Nevertheless, the fact that *yolk-GAL4* drives the highest transgene transcriptional activity in the fat body suggests that there might be a threshold level of GAL4 production past which adipocyte functional capacities are overwhelmed, triggering major fat body tissue failure. Additionally, specific molecular properties of GAL4 protein might exert cytotoxic effects that are most pronounced at the levels observed only in *yolk-GAL4* females. Our observation that cytoplasmic and nuclear-localized mCherry, and LexA protein had a less severe impact on infection survival (Fig.6H, Fig.7G, Fig.S8D, Fig.S9E) and did not recapitulate the energy storage and reproductive phenotypes of *yolk-GAL4* (Fig.6E,F,G,

Fig.7D-F, Fig.S8A,B,C, Fig.S9B,C,D) would be consistent with a specific toxicity of GAL4. Because the LexA::GAD fusion protein includes the GAL4 activation domain, our results suggest that whatever attributes of GAL4 might specifically induce toxicity are likely to involve the C-terminal DNA binding domain or properties only conferred by the full protein conformation. The potential for high levels of GAL4 to trigger major tissue stress in the fat body by overwhelming core cellular processes should be explored further, particularly as this might yield fundamental insights into how adipocytes and other cell types allocate resources among competing biological processes and the limits of their molecular functional capacities.

In addition to general cellular strain, we also considered the possibility that GAL4 might induce apoptosis of fat body cells, as similar effects were documented in two prior studies. The *GMR-GAL4* driver induced programmed cell death in the posterior region of eye imaginal discs (Kramer and Staveley 2003; Rezával et al. 2007), leading to degenerated ommatidia in adult eyes, while *pdf-GAL4* caused apoptosis of circadian neurons, leading to disrupted sleep-wake behaviors (Rezával et al. 2007). In both of these cases, the phenotypic consequences of GAL4-induced cell death could be partially suppressed by expressing the apoptosis-inhibiting p35 transgene (Rezával et al. 2007). Similarly, Kim et al. (2014), showed that p35 moderately restored infection-induced *Diptericin* expression in *yolk-GAL4* heterozygotes. In contrast to these reports, we found that driving p35 did not suppress the reduced fat stores, low egg output, or sensitivity to *P. rettgeri* infection of flies expressing *yolk-GAL4* (Fig.S6). Importantly, however, these data do not fully exclude the possibility that GAL4 might induce programmed death of fat body cells. For example, GAL4 might trigger apoptosis via the initiator caspase Dronc, which is not affected by p35 (Hawkins et al. 2000; Meier et al. 2000). Beyond apoptotic effects, additional evidence suggests GAL4 can disrupt cellular protein homeostasis and quality control mechanisms. In a microarray analysis of GAL4-induced gene expression changes in larval salivary glands, Liu and Lehmann (2008) identified protein ubiquitination factors as overrepresented among the GAL4-responsive genes. The muscle-specific MHC-GeneSwitchGAL4 transgene was recently shown to cause age-dependent accumulation of polyubiquitinated protein aggregates in adult flight muscles, although these effects required activation of the GAL4-progesterone receptor fusion by RU486 feeding (Zappia et al. 2024). Similarly, Rezaval et al. (2007) found high levels of GAL4 in insoluble protein extracts from heads of aged flies expressing *pdf-GAL4*. Collectively, these reports suggest that GAL4 protein itself might form insoluble aggregates, particularly when expressed at high levels. In the adult fat body, accumulation of these protein aggregates could disrupt the function of organelles necessary for lipid anabolism and yolk export and could trigger tissue loss via alternative routes like necrotic cell death and autophagic clearance. Future work should consider the scope of potential mechanisms by which GAL4 might compromise adipocyte function and viability.

We found that three distinct nuclear-localized transgenic proteins–GAL4, mCherry, and LexA–reduced flies’ capacity to survive systemic bacterial infection, with severity dependent on the level of transgene expression in the fat body or on transgene copy number (Fig.4, Fig.5E,F, Fig.6H, Fig.7G, Fig.S5B, Fig.S8D, Fig.S9E). Importantly, we observed that the identical mCherry protein expressed from the same genomic locus and in the same tissue-specific pattern but lacking nuclear localization sequence had no effect on infection mortality (Fig.6H). This specifically implicates the process of nuclear import and/or presence of transgenic proteins in fat body nuclei as major determinants of infection sensitivity. In *Drosophila* and other insects, systemic microbial infections activate a broadly conserved humoral immune response that is predominantly regulated by the Immune Deficiency (IMD) and Toll signaling pathways. These pathways comprise intracellular signaling cascades that culminate in the nuclear translocation of the NF-κB family transcription factors Relish (Rel), Dorsal-related immunity factor (Dif), and Dorsal (Dl). On entering the nucleus, these transcription factors massively upregulate the expression of AMPs that are then secreted from the fat body into the hemolymph to kill the pathogen (De Gregorio et al. 2002; Valanne et al. 2011; Myllymäki et al. 2014; Troha & Im et al. 2018; Westlake et al. 2025). In the larval fat body, a specific nucleoporin encoded by the gene *members only* (*mbo*, homologous to vertebrate *Nup88*) is required for nuclear import of Rel, Dif, and Dl upon infection (Uv et al. 2000). Interestingly, *mbo* is also required for nuclear localization of GAL4 (Uv et al. 2000). Our data are consistent with a model where high levels of nuclear-localized transgenic proteins might oversaturate nuclear pore channels, such as those formed with mbo, and physically impede the infection-induced nuclear translocation of NF-κBs resulting in abrogated or delayed AMP production and consequent failure to constrain pathogen proliferation. In further support of this model, we and Kim et al. (2014) found that infected flies expressing *yolk-GAL4* display markedly reduced AMP gene upregulation (Fig.4I), which could reflect impaired nuclear import of Rel.

Along with effects on nuclear import processes, accumulation of intranuclear GAL4 could also affect nuclear and chromatin architecture through non-specific interactions with structural proteins like lamins or DNA organizational factors like histones. This structural disruption might physically hinder transcription factors from efficiently accessing the regulatory sequences of their target genes. The potential for GAL4 and other nuclear-localized transgenic proteins to affect nuclear import and function of native transcription factors warrants further investigation, particularly as this could affect studies of other physiological processes that depend on the rapid induction of transcriptional programs, such as starvation (Wang et al. 2008), heavy metal stress (Yepiskoposyan et al. 2006), and oxidative stress responses (Sykiotis and Bohmann 2008). Additionally, if this model is correct, the exact molecular properties and interactions through which GAL4 and other transgenic proteins affect nuclear import processes could be leveraged to develop experimental tools such as peptide agonists to selectively perturb the subcellular localization and function of specific transcription factors. Such tools could enable detailed investigation of the nuclear shuttling kinetics and threshold levels required for endogenous transcription factors to elicit effects on gene expression and physiology.

In conducting this study, we generated several new *D. melanogaster* lines that will be broadly useful to the *Drosophila* research community. Our *yolk-mCherry* lines faithfully recapitulate the well-documented spatiotemporal and sex-specific expression patterns of the *yolk* enhancer (Fig.S1, Fig.S7), and effectively re-create historical *yolk-lacZ* reporter lines that yielded fundamental discoveries about mechanisms of enhancer regulation (Coschigano and Wensink 1993; An and Wensink 1995a; An and Wensink 1995b) but, to our knowledge, are now lost. Both *yolk-mCherry.nls* and *yolk-mCherry.cyt* provide robust, easily-visualized fluorescent readouts of *yolk* enhancer activity, and could be used as indicators for studying processes such as sex determination (Belote et al. 1985; Bownes 1994; Tarone et al. 2012), nutrient sensing systems (Bownes and Blair 1986; Bownes et al. 1988; Terashima et al. 2005), and endocrine signaling (Jowett and Postlethwait 1980; Bownes et al. 1987; Bownes et al. 1996). In addition to these reporters, we used HACK (Lin and Potter 2016; Chang et al. 2022; Rankin et al. 2024) to convert *yolk-GAL4* into *yolk-LexA*, which can be used in genetic experiments employing the *LexA/LexAop* binary expression system (Lai and Lee 2006). Crucially, and in striking contrast to *yolk-GAL4*, heterozygosity for *yolk-LexA* (as would be the case in most experimental setups) did not reduce fat levels or egg production (Fig.7D-F, Fig.S9B,C,D), and little or no effect on sensitivity to infection (Fig.7G, Fig.S9E). We have therefore generated a reagent that retains the major advantages of inherent adult female-specific expression conferred by the *yolk* enhancer, but which lacks the extreme deleterious effects on fly physiology caused by *yolk-GAL4*. As such, the *yolk-LexA* line constitutes a valuable new tool facilitating genetic manipulations to investigate adult fat body functions.

Given the central utility of the *GAL4-UAS* system for answering fundamental biological questions with *Drosophila*, our findings emphasize the importance of thoroughly characterizing individual driver lines and thoughtful selecting genetic reagents when designing experiments. For example, a highly-expressed GAL4 driver will likely achieve more efficient RNAi knockdown of a gene of interest, but resulting phenotypes could be confounded by the driver’s deleterious impacts on the target cell type or tissue. On the other hand, a weaker driver will likely affect tissue function to a lesser degree but might not activate expression of the RNAi hairpin construct strongly enough to disrupt the function of the gene being studied. Ideally, when possible, drivers should be selected that strike a balance between these two scenarios, activating *UAS* transgenes sufficiently to achieve the intended experimental manipulation but without causing overt defects in the focal tissue. In all experiments, flies expressing transgenic components alone should compared to non-transgenic flies to evaluate how *GAL4* (or other relevant transgenes) might affect the trait being assayed.

Beyond *Drosophila*, our study contributes to a body of work cautioning potential inadvertent effects of heterologous protein expression in varied biological systems. In *Saccharomyces cerevisiae*, strong expression of eGFP and other fluorescent reporter proteins disrupts proteasomal function and affects cell morphology and growth (Kafri et al. 2016; Kintaka et al. 2016; Kintaka et al. 2020; Namba et al. 2022; Fujita et al. 2024; Namba and Moriya 2024). Similarly, GFP expressed in cultured cell lines and rodent models can induce apoptosis and elicit deleterious oxidative stress and immunological responses (Liu et al. 1999; Huang et al. 2000; Detrait et al. 2002; Krestel et al. 2004; Agbulut et al. 2006; Agbulut et al. 2007; Ansari et al. 2016; Kalyanaraman and Zielonka 2017; Verma et al. 2024). Apoptotic neuronal cell death induced by co-expression of eGFP and Δ-galactosidase in mouse forebrains has been shown to impair motor coordination and cause rapid, early mortality (Krestel et al. 2004). Verma et al. (2024) recently reported severe defects in mouse eye development caused by expression of a Tet-inducible, nuclear-localized GFP reporter construct. Notably, in multiple cases these studies found that higher GFP expression levels, resulting from homozygosity for the transgene or increasing promoter strength, resulted in greater phenotype severity. This aligns with our observations of dose-dependent effects of transgene expression in the *Drosophila* fat body. These prior reports in diverse systems combined with our present work underscore the fact that transgenic proteins employed as experimental “tools” occur within complex, functioning cellular environments, and therefore have the potential to directly or indirectly impact many cellular components and processes. These effects should be empirically evaluated and taken into consideration when designing and interpreting experiments employing the *GAL4-UAS* system and other transgenic systems.

## DATA AVAILABILITY

*Drosophila* lines and plasmids are available upon request. The authors affirm that all data necessary for confirming the conclusions of the article are present within the article, figures, and tables.

## ACKNOWLEDGMENTS

We are grateful to members of the Lazzaro lab for helpful discussions and advice throughout the completion of this study. We thank Kate Browning for technical assistance with capturing stereoscope images of adult fat bodies. We thank Justin DiAngelo (Penn State Berks) and Alissa Armstrong (University of South Carolina) for sharing fly stocks. We thank Sangbin Park and Seung Kim (Stanford University) for sharing *CyO, LexA-HACKy+* donor flies and providing advice on the HACK approach. Stocks obtained from the Bloomington *Drosophila* Stock Center (NIH P40OD018537) were used in this study.

## SUPPLEMENTARY TABLES

**Table S1.**
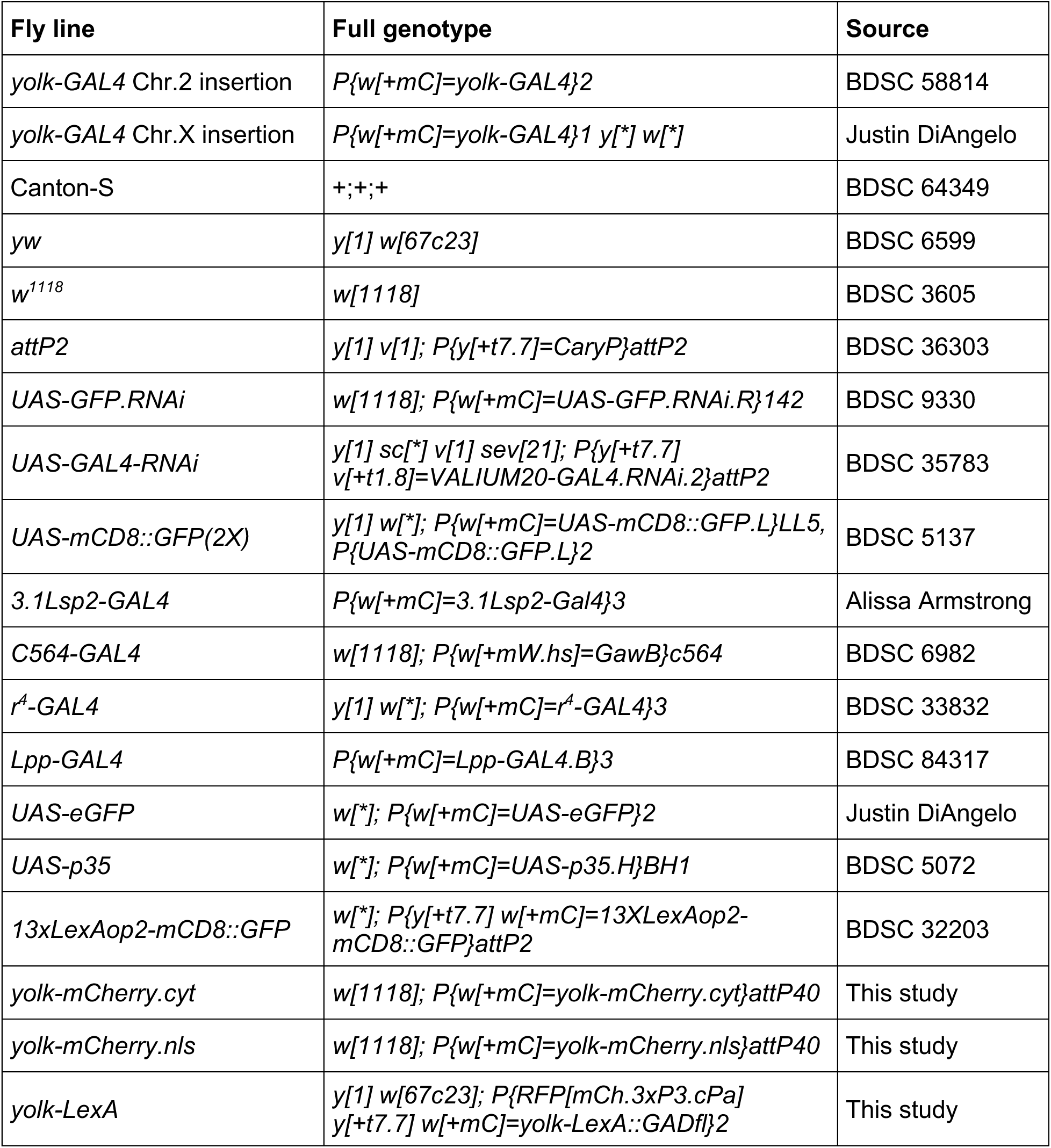
Fly lines used in this study.

**Table S2.**
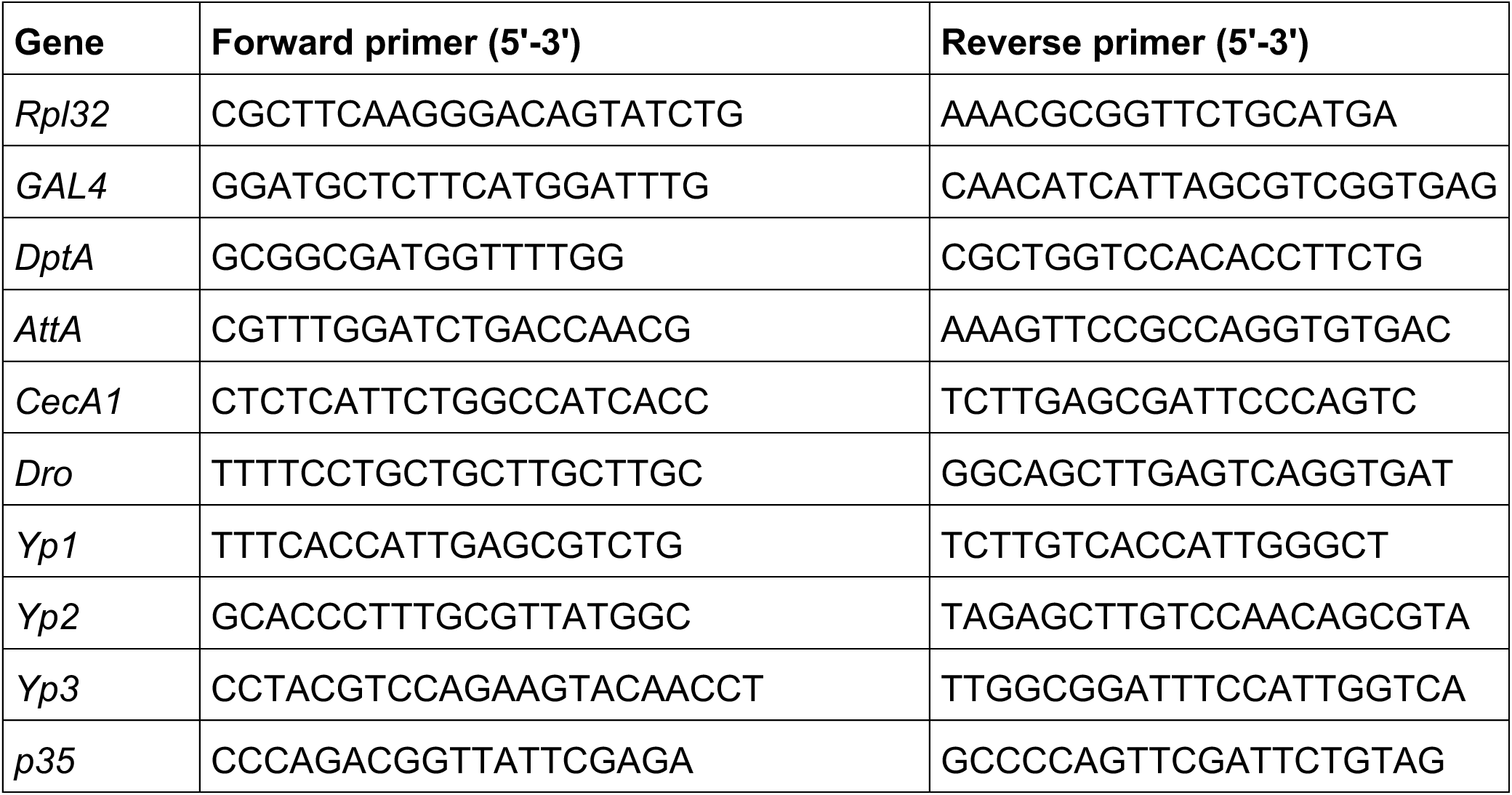
Primers used for RT-qPCR and RT-PCR.

**Table S3.** Sample sizes for data represented in all figure panels.

| Figure | Genotype/condition | Assay | n (# replicates) | n (# individuals) |
| --- | --- | --- | --- | --- |
| 1F | P{yolk-GAL4}2/+ (attP2) | GAL4 expression | 4 | 5-10 pooled flies per replicate |
|  | P{yolk-GAL4}2>GAL4-RNAi |  | 5 |  |
|  | P{yolk-GAL4}1/+ (attP2) |  | 6 |  |
|  | P{yolk-GAL4}1>GAL4-RNAi |  | 6 |  |
| 2B | Canton-S | ORO signal intensity | 3 | 6 |
|  | P{yolk-GAL4}2/+ |  | 3 | 5 |
|  | P{yolk-GAL4}2/P{yolk-GAL4}2 |  | 3 | 6 |
| 2C | Canton-S | Starvation resistance | 3 | 88 |
|  | P{yolk-GAL4}2/+ |  | 3 | 100 |
|  | P{yolk-GAL4}2/P{yolk-GAL4}2 |  | 3 | 94 |
| 2E | P{yolk-GAL4}2/+ (attP2) | ORO signal intensity | 3 | 5 |
|  | P{yolk-GAL4}2>GFP-RNAi |  | 3 | 6 |
|  | P{yolk-GAL4}2>GAL4-RNAi |  | 3 | 5 |
| 2F | P{yolk-GAL4}2/+ (attP2) | Starvation resistance | 3 | 114 |
|  | P{yolk-GAL4}2>GFP-RNAi |  | 3 | 102 |
|  | P{yolk-GAL4}2>GAL4-RNAi |  | 3 | 139 |
| S2B | yw | ORO signal intensity | 3 | 8 |
|  | P{yolk-GAL4}1/+ |  | 3 | 6 |
|  | P{yolk-GAL4}1/P{yolk-GAL4}1 |  | 3 | 7 |
| S2C | yw | Starvation resistance | 3 | 70 |
|  | P{yolk-GAL4}1/+ |  | 3 | 119 |
|  | P{yolk-GAL4}1/P{yolk-GAL4}1 |  | 3 | 108 |
| S2E | P{yolk-GAL4}1/+ (attP2) | ORO signal intensity | 3 | 8 |
|  | P{yolk-GAL4}1>GFP-RNAi |  | 3 | 5 |
|  | P{yolk-GAL4}1>GAL4-RNAi |  | 3 | 7 |
| S2F | P{yolk-GAL4}1/+ (attP2) | Starvation resistance | 3 | 129 |
|  | P{yolk-GAL4}1>GFP-RNAi |  | 3 | 125 |
|  | P{yolk-GAL4}1>GAL4-RNAi |  | 3 | 123 |
| 3A | Canton-S | Egg laying | 3 | 9 |
|  | P{yolk-GAL4}2/+ |  | 3 | 8 |
|  | P{yolk-GAL4}2/P{yolk-GAL4}2 |  | 3 | 9 |
| 3B | Canton-S | Ovary size | 3 | 17 |
|  | P{yolk-GAL4}2/+ |  | 3 | 14 |
|  | P{yolk-GAL4}2/P{yolk-GAL4}2 |  | 3 | 14 |
| 3D | P{yolk-GAL4}2/+ (attP2) | Egg laying | 3 | 9 |
|  | P{yolk-GAL4}2>GFP-RNAi |  | 3 | 9 |
|  | P{yolk-GAL4}2>GAL4-RNAi |  | 3 | 8 |
| 3E | P{yolk-GAL4}2/+ (attP2) | Ovary size | 3 | 14 |
|  | P{yolk-GAL4}2>GFP-RNAi |  | 3 | 18 |
|  | P{yolk-GAL4}2>GAL4-RNAi |  | 3 | 13 |
| 3G | Canton-S | Yolk Protein gene expression | 4 | 10 pooled fat bodies/abdominal carcasses per replicate |
|  | P{yolk-GAL4}2/+ |  | 4 |  |
| 3I | Canton-S | # Vitellogenic follicles | 3 | 18 |
|  | P{yolk-GAL4}2/+ |  | 3 | 16 |
| 3J | Canton-S | # Mature oocytes | 3 | 45 |
|  | P{yolk-GAL4}2/+ |  | 3 | 52 |
| S3A | yw | Egg laying | 5 | 11 |
|  | P{yolk-GAL4}1/+ |  | 5 | 7 |
|  | P{yolk-GAL4}1/P{yolk-GAL4}1 |  | 3 | 9 |
| S3B | yw | Ovary size | 3 | 12 |
|  | P{yolk-GAL4}1/+ |  | 3 | 18 |
|  | P{yolk-GAL4}1/P{yolk-GAL4}1 |  | 3 | 14 |
| S3D | P{yolk-GAL4}1/+ (attP2) | Egg laying | 5 | 10 |
|  | P{yolk-GAL4}1>GFP-RNAi |  | 5 | 10 |
|  | P{yolk-GAL4}1>GAL4-RNAi |  | 4 | 8 |
| S3E | P{yolk-GAL4}1/+ (attP2) | Ovary size | 3 | 17 |
|  | P{yolk-GAL4}1>GFP-RNAi |  | 3 | 15 |
|  | P{yolk-GAL4}1>GAL4-RNAi |  | 3 | 17 |
| S3I | P{yolk-GAL4}2/+ (attP2) | GAL4 expression | 4 | 10 pooled ovaries per replicate |
|  | P{yolk-GAL4}1/+ (attP2) |  | 4 |  |
| S3J | P{yolk-GAL4}2/+ | Embryo hatch rate | 9 | 180 |
|  | P{yolk-GAL4}2/P{yolk-GAL4}2 |  | 7 | 141 |
| 4A | Canton-S | P. rettgeri infection survival | 3 | 107 |
|  | P{yolk-GAL4}2/+ |  | 3 | 80 |
|  | P{yolk-GAL4}2/P{yolk-GAL4}2 |  | 3 | 72 |
| 4B | Canton-S | E. faecalis infection survival | 3 | 104 |
|  | P{yolk-GAL4}2/+ |  | 3 | 79 |
|  | P{yolk-GAL4}2/P{yolk-GAL4}2 |  | 3 | 66 |
| 4C | Canton-S | Ecc15 infection survival | 3 | 108 |
|  | P{yolk-GAL4}2/+ |  | 3 | 149 |
|  | P{yolk-GAL4}2/P{yolk-GAL4}2 |  | 3 | 148 |
| 4D | yw | P. rettgeri infection survival | 3 | 62 |
|  | P{yolk-GAL4}1/+ |  | 3 | 80 |
|  | P{yolk-GAL4}1/P{yolk-GAL4}1 |  | 3 | 77 |
| 4E | P{yolk-GAL4}2/P{yolk-GAL4}2 males | P. rettgeri infection survival | 3 | 73 |
|  | P{yolk-GAL4}2/P{yolk-GAL4}2 females |  | 3 | 68 |
| 4F | P{yolk-GAL4}1/P{yolk-GAL4}1 males | P. rettgeri infection survival | 3 | 87 |
|  | P{yolk-GAL4}1/P{yolk-GAL4}1 females |  | 3 | 86 |
| 4G | P{yolk-GAL4}2/+ (attP2) | P. rettgeri infection survival | 3 | 143 |
|  | P{yolk-GAL4}2>GFP-RNAi |  | 3 | 90 |
|  | P{yolk-GAL4}2>GAL4-RNAi |  | 3 | 166 |
| 4H | P{yolk-GAL4}1/+ (attP2) |  | 3 | 68 |
|  | P{yolk-GAL4}1>GFP-RNAi |  | 3 | 81 |
|  | P{yolk-GAL4}1>GAL4-RNAi | P. rettgeri infection survival | 3 | 82 |
| 4I | Canton-S uninfected | AMP expression | 4 | 10 pooled fat bodies/abdominal carcasses per replicate |
|  | Canton-S P. rettgeri infected |  | 4 |  |
|  | P{yolk-GAL4}2/+ uninfected |  | 4 |  |
|  | P{yolk-GAL4}2/+ P. rettgeri infected |  | 4 |  |
| 4J | Canton-S 0h | P. rettgeri bacterial loads | 3 | 19 |
|  | P{yolk-GAL4}2/+ 0h |  | 3 | 17 |
|  | Canton-S 6h |  | 3 | 27 |
|  | P{yolk-GAL4}2/+ 6h |  | 3 | 28 |
|  | Canton-S 12h |  | 3 | 27 |
|  | P{yolk-GAL4}2/+ 12h |  | 3 | 26 |
|  | Canton-S 18h |  | 3 | 27 |
|  | P{yolk-GAL4}2/+ 18h |  | 3 | 30 |
| 5A | 3.1Lsp2-GAL4/+ | GAL4 expression | 3 | 10 pooled fat bodies/abdominal carcasses per replicate |
|  | C564-GAL4/+ |  | 4 |  |
|  | r4-GAL4/+ |  | 4 |  |
|  | Lpp-GAL4/+ |  | 4 |  |
|  | P{yolk-GAL4}2/+ |  | 4 |  |
| 5C | 3.1Lsp2-GAL4/+ | ORO signal intensity | 3 | 10 |
|  | C564-GAL4/+ |  | 3 | 7 |
|  | r4-GAL4/+ |  | 3 | 10 |
|  | Lpp-GAL4/+ |  | 3 | 9 |
|  | P{yolk-GAL4}2/+ |  | 3 | 7 |
| 5D | 3.1Lsp2-GAL4/+ | Egg laying | 8 | 18 |
|  | C564-GAL4/+ |  | 7 | 17 |
|  | r4-GAL4/+ |  | 8 | 22 |
|  | Lpp-GAL4/+ |  | 7 | 16 |
|  | P{yolk-GAL4}2/+ |  | 7 | 20 |
| 5E | 3.1Lsp2-GAL4/+ | P. rettgeri infection survival | 5 | 100 |
|  | C564-GAL4/+ |  | 5 | 164 |
|  | r4-GAL4/+ |  | 5 | 148 |
|  | Lpp-GAL4/+ |  | 5 | 99 |
|  | P{yolk-GAL4}2/+ |  | 5 | 111 |
| S5A | Lpp-GAL4/+ (attP2) | GAL4 expression | 3 | 5-10 pooled flies per replicate |
|  | Lpp-GAL4>GAL4-RNAi |  | 3 |  |
| S5B | Lpp-GAL4/+ (attP2) | P. rettgeri infection survival | 3 |  |
|  | Lpp-GAL4>GFP-RNAi |  | 3 | 63 |
|  | Lpp-GAL4>GAL4-RNAi |  | 3 | 64 |
| S6C | P{yolk-GAL4}2>GFP | ORO signal intensity | 3 | 8 |
|  | P{yolk-GAL4}2>p35 |  | 3 | 10 |
| S6D | P{yolk-GAL4}2>GFP | Egg laying | 6 | 16 |
|  | P{yolk-GAL4}2>p35 |  | 4 | 10 |
| S6E | P{yolk-GAL4}2>GFP | P. rettgeri infection survival | 3 | 93 |
|  | P{yolk-GAL4}2>p35 |  | 3 | 78 |
| 6F | w1118 | ORO signal intensity | 3 | 10 |
|  | yolk-mCherry.nls/+ |  | 3 | 11 |
|  | yolk-mCherry.nls/yolk-mCherry.nls |  | 3 | 8 |
|  | yolk-mCherry.cyt/+ |  | 3 | 7 |
|  | yolk-mCherry.cyt/yolk-mCherry.cyt |  | 3 | 9 |
| 6G | w1118 | Egg laying | 5 | 13 |
|  | yolk-mCherry.nls/+ |  | 5 | 13 |
|  | yolk-mCherry.nls/yolk-mCherry.nls |  | 5 | 11 |
|  | yolk-mCherry.cyt/+ |  | 5 | 12 |
|  | yolk-mCherry.cyt/yolk-mCherry.cyt |  | 5 | 12 |
| 6H | w1118 | P. rettgeri infection survival | 4 | 99 |
|  | yolk-mCherry.nls/+ |  | 4 | 87 |
|  | yolk-mCherry.nls/yolk-mCherry.nls |  | 4 | 89 |
|  | yolk-mCherry.cyt/+ |  | 4 | 101 |
|  | yolk-mCherry.cyt/yolk-mCherry.cyt |  | 4 | 77 |
| 7E | yw | ORO signal intensity | 3 | 10 |
|  | yolk-LexA/+ |  | 3 | 6 |
|  | yolk-LexA/yolk-LexA |  | 3 | 10 |
| 7F | yw | Egg laying | 7 | 14 |
|  | yolk-LexA/+ |  | 8 | 16 |
|  | yolk-LexA/yolk-LexA |  | 8 | 15 |
| 7G | yw | P. rettgeri infection survival | 5 | 96 |
|  | yolk-LexA/+ |  | 5 | 73 |
|  | yolk-LexA/yolk-LexA |  | 5 | 112 |
| S8B | w1118/+;+/+ | ORO signal intensity | 3 | 8 |
|  | w1118/+;yolk-mCherry.nls/+ |  | 3 | 9 |
|  | w1118/+;yolk-mCherry.cyt/+ |  | 3 | 9 |
| S8C | w1118/+;+/+ | Egg laying | 6 | 18 |
|  | w1118/+;yolk-mCherry.nls/+ |  | 6 | 16 |
|  | w1118/+;yolk-mCherry.cyt/+ |  | 6 | 15 |
| S8D | w1118/+;+/+ | P. rettgeri infection survival | 4 | 120 |
|  | w1118/+;yolk-mCherry.nls/+ |  | 4 | 70 |
|  | w1118/+;yolk-mCherry.cyt/+ |  | 4 | 98 |
|  | P{yolk-GAL4}2/+ |  | 4 | 50 |
| S9C | yw/+;+/+ | ORO signal intensity | 3 | 8 |
|  | yw/+;yolk-LexA/+ |  | 3 | 10 |
| S9D | yw/+;+/+ | Egg laying | 5 | 10 |
|  | yw/+;yolk-LexA/+ |  | 5 | 10 |
| S9E | yw/+;+/+ | P. rettgeri infection survival | 4 | 99 |
|  | yw/+;yolk-LexA/+ |  | 4 | 105 |
|  | P{yolk-GAL4}2/+ |  | 4 | 53 |

